# The RNA Editome of Esophageal Squamous Cell Carcinoma Identifies an ADAR1-CDK13 Editing Feedback Loop Mediating cGAS–STING Activation in Early Tumorigenesis

**DOI:** 10.64898/2026.01.29.701210

**Authors:** Lihong Huang, Minglei Yang, Douyue Li, Guozhong Jiang, Wenliang Zhang

**Affiliations:** The Key Laboratory of Advanced Interdisciplinary Studies, The First Affiliated Hospital of Guangzhou Medical University, GMU-GIBH Joint School of Life Sciences, Guangdong Provincial Key Laboratory of Protein Modification and Disease, The Guangdong-Hong Kong-Macao Joint Laboratory for Cell Fate Regulation and Diseases, Guangzhou Medical University, Guangzhou 510182, China; Department of Pathology, The First Affiliated Hospital of Zhengzhou University, Zhengzhou 450000, Henan, China; Department of Hepatobiliary Surgery, The First Affiliated Hospital of Guangzhou Medical University, Guangzhou 510120, China

**Keywords:** RNA editing, *CDK13*, *ADAR1*, Esophageal Squamous Cell Carcinoma, cGAS–STING, Interferon-stimulated genes

## Abstract

**Objective:** RNA adenosine-to-inosine (A-to-I) editing, catalyzed by adenosine deaminases acting on RNA (ADARs), is a pervasive post-transcriptional mechanism that expands transcriptomic and proteomic diversity. However, the global landscape of RNA A-to-I editing, as well as its functional and clinical significance in esophageal squamous cell carcinoma (ESCC), remains largely unexplored. This study aimed to systematically characterize the RNA editome of ESCC and elucidate its biological and clinical significance.

**Methods:** Whole-transcriptome sequencing was performed on 121 primary ESCC tumors, with or without lymph node metastasis, together with matched normal tissues, to construct a high-resolution RNA editome. ADAR1-regulated RNA editing events were identified, and their functional consequences were investigated using integrated transcriptomic, phosphoproteomic, and RNA immunoprecipitation sequencing (RIP-seq) analyses. Associations between *CDK13* editing, cGAS–STING–interferon-stimulated gene (ISG) signaling, and patient survival were further evaluated.

**Results:** A total of 222,020 high-confidence RNA editing sites were identified, of which approximately 98% were A-to-I events, including 124,486 ESCC-specific edits predominantly enriched in introns, 3′ untranslated regions, and non-coding RNAs, highlighting a pervasive post-transcriptional regulatory layer. RNA A-to-I editing was extensively remodeled and globally up-regulated in non-metastatic ESCC, whereas only minimal changes were observed during lymph node metastasis, indicating that RNA editing alterations predominantly occur during early tumorigenesis. *CDK13* emerged as a recurrent ADAR1 target, with RNA editing inversely correlated with *CDK13* expression. ADAR1-mediated *CDK13* editing established a positive feedback loop associated with enhanced interferon-stimulated gene (ISG) signaling and poorer survival in non-metastatic ESCC. Mechanistically, partial attenuation of *CDK13* induced chronic, tumor cell–intrinsic activation of the cGAS–STING–ISG pathway. Integrated multi-omics analyses further revealed that *CDK13* regulates phosphorylation networks governing cytoskeleton organization, intracellular trafficking, RNA homeostasis, and immune signaling.

**Conclusion:** RNA A-to-I editing represents a dynamic regulatory mechanism driving early ESCC progression and remodeling tumor cell–intrinsic immune signaling. ADAR1-mediated editing of *CDK13* provides a mechanistic link between RNA editing and cGAS–STING–ISG pathway activation, revealing potential therapeutic vulnerabilities and supporting its utility as an early prognostic biomarker in ESCC.

## Introduction

Esophageal cancer, ranking sixth of among all major cancer diseases, possesses high incidence and mortality rates worldwide [1]. It consists of two main histological subtypes: adenocarcinoma (EAC) and squamous cell carcinoma (ESCC), which differ in etiological and molecular features [2]. EAC is associated with obesity and gastroesophageal reflux disease (GERD), while ESCC is linked to abusive smoking and alcohol consumption. Despite the advances in traditional therapy, the survival rate of esophageal cancer remains low, underscoring the urgent need for novel biomarkers and RNA-based therapeutic strategies.

RNA editing, as one of the most prevalent post-transcriptional RNA modifications, can alter nucleotide sequence of RNA species, leading to transcriptome and proteome diversity and complexity. It is catalyzed by enzymes under specific regulation involving mainly adenosine to inosine (A-to-I) and cytidine to uridine (C-to-U) conversions. In cDNA sequence, A-to-G editing is considered A-to-I editing because of the reading of inosine to guanidine by reverse transcriptase. Vertebrates contain three ADAR proteins: ADAR1 (ADAR, with two main isoforms p110 and p150), ADAR2 (ADARB1), and ADAR3 (ADARB2). ADAR1 and ADAR2 as two major editors are ubiquitously expressed, whereas ADAR3 is dominant in the brain and function as negative regulator due to lack of deaminase activity[3]. ADARs are crucial for developmental processes. Heterozygous *Adar1^+/-^* in mice embryo caused lethality [4, 5]. *adar-1* and *adar-2* double knockout mutants in *C. elegans* defected in chemotaxis [6]. Mice embryo containing editing deficient knock-in mutation *Adar1^E861A/E861A^* died in early embryonic day, while this phenotype was rescued when concurrently deleted dsRNA sensor, MDA5 [7].

RNA editing primarily targets within double-stranded regions of distinct RNA species. Numerous A-to-I cDNA changes were identified that originated from diverse genomic regions such as introns and 3′ untranslated regions (UTRs) through deep sequencing. Reports showed that several transcripts undergo RNA editing in cancer, affecting their expression and activity [8]. For instance, the conformation of the amino acid sequence of serotonin 2C receptor (5-HT_2c_R) (5HT, 5-hytroxytryptamine) is changed by editing on adenosine residue [9], resulting in a significant reduction of conjugation efficiency between 5-HT_2C_R and G protein [9–13]. Another well-known editing on pre-mRNA is α-amino-3-hydroxy-5-methyl-isoxazole-4-propionic acid (AMPA) [14–17] and kainate receptor [18]. An interrupted nonsynonymous mutation on the R/G site catalyzed by ADAR2 leads to Amyotrophic Lateral Sclerosis (ALS) [17, 19–23], indicating the essential role of RNA editing in human diseases. In addition, blocking A-to-I editing of *AZIN1* transcripts by ADAR1 may change protein conformation and attenuates translocation in hepatocellular carcinoma [24]. Editing in *GLI1* at position 701 that leads to nonsynonymous mutation impaired *GLI1* transcription and *GLI1* related cellular proliferation [25] and advanced malignant regeneration in myeloma [26]. Beyond pre-mRNAs, RNA editing intersects with RNA interference pathways, such as in C. elegans [27], and can affect miRNA processing and target regulation, exemplified by editing of the miR-376 cluster [28] and the 3′ UTR of MDM2 [29]. Despite these advances, the landscape of RNA A-to-I editing and the mechanisms of ADAR1-dependent editing in esophageal squamous cell carcinoma (ESCC) remain largely unexplored. Recent studies indicate that ADAR enzymes play critical roles in ESCC: ADAR1 is overexpressed in sporadic ESCC and drives hyperediting of oncogenic transcripts such as *AZIN1*, contributing to aggressive tumor behavior [30]; ADAR2 mediates RNA editing of *SLC22A3* in familial ESCC, promoting early invasion and metastasis [31]; and ADAR proteins can also exert RNA editing-independent oncogenic functions, such as stabilizing the deubiquitinase USP38 [32]. Although individual edited transcripts such as *AZIN1* and *SLC22A3* have been implicated in tumor progression, the global landscape of RNA A-to-I editing, its functional consequences, and its potential role in ESCC remain poorly understood.

To address this gap, we performed whole-transcriptome sequencing of 121 primary ESCC tumors, with or without lymph node metastasis, together with matched normal tissues, to systematically map RNA A-to-I editing events. By integrating transcriptomic profiling with ADAR1 RNA immunoprecipitation sequencing (RIP-seq), we identified widespread ADAR1-catalyzed editing, predominantly affecting introns, 3′-untranslated regions, and non-coding RNAs. A-to-I editing was extensively altered and largely upregulated in non-metastatic ESCC, with minimal changes during lymph node metastasis. Among recurrently edited genes, *CDK13* emerged as a key ADAR1 target, who’s editing forms a positive feedback loop that drives interferon-stimulated gene (ISG) signaling by remodeling phosphorylation networks that regulate cytoskeletal organization, intracellular trafficking, RNA homeostasis, and cGAS–STING signaling. These findings indicate that ADAR1-mediated RNA editing reshapes innate immune pathways through tumor cell – intrinsic cGAS – STING activation, thereby contributing to early ESCC tumorigenesis.

## Results

### High-confidence RNA editing landscape in ESCC and matched normal tissues

We employed whole-transcriptome sequencing with a paired-end library featuring a read length of 2 × 150 bp yielding a total of 100-125 million reads per sample (5-6.2 × coverage; **Supplementary Table 1**), which provided a global genomic landscape for esophageal squamous cell carcinoma (ESCC) and matched adjacent normal tissues (CTRL). Poisson distance heatmap and MDA plot demonstrated the clear separation of transcriptome profile of ESCC and CTRL samples (**Supplementary Fig. S1**). This approach surpasses previous RNA editing studies involved in mRNA-seq with shorter sequencing lengths and improves the resolution to detect editing events [33–35].

To achieve high-quality RNA editing analysis, we leveraged combined tools to conduct pre-processing before calling with pileup and calls (**Fig. 1a**), as it improved the accuracy of identification and bcftools mpileup has better performance than GATK Haplotypecaller [36]. Specifically, RNA-seq reads were aligned to the GRCh38 reference genome, followed by hard filtering to remove low-quality reads and those mapped to ribosomal RNAs (rRNAs). The resulting data was subjected to pre-processed using the GATK toolkits and RNA-DNA mismatches were identified using bcftools (**Fig. 1b**), followed by rigorous filtering to remove known DNA mutations (**Fig. 1c**), low-confidence calls (**Fig. 1d**), and sites proximal to splice junctions and with low prevalence among patients (n = 6) (**Fig. 1e**). Most identified sites corresponded to canonical A-to-I editing events, which remained dominant throughout all filtering steps.

**Fig. 1.**
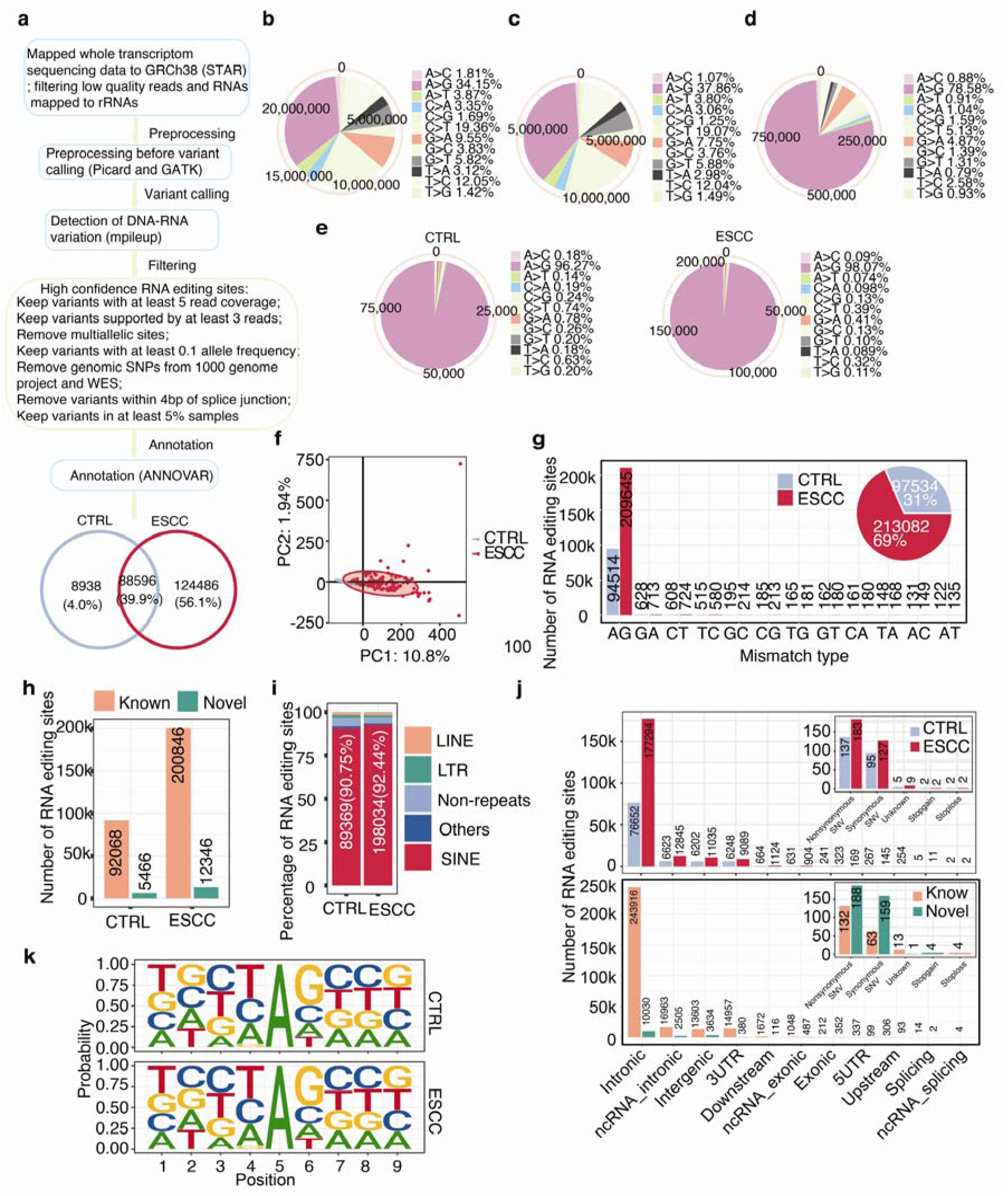
Identification of high-confidence RNA editing mismatches with the combined method. **(a)** Pipeline illustrating pre-processing, calling, and filtering for RNA editing sites. **(b)** Variants called in ESCC and matched normal samples (CTRL). **(c)** Variants retained after removal of multiallelic sites and known DNA SNVs (dbSNP138 & 1000genome & WES). **(d)** Variants retained by keeping mismatches with QUAL above 30 and read depth more than 10 and supported by at least 3 reads with at least 0.1 allele frequencies. **(e)** Variants retained in ESCC and CTRL by keeping mismatches within 4bp of the splice junction and in at least 5% samples. **(f)** PCA plot showing variation of RNA A-to-I editing across samples in ESCC and CTRL cohort. Mismatches present in more than 50% samples were retained for the analysis. **(g)** RNA mismatched types in ESCC and CTRL. **(h)** Abundance of known and novel RNA editing sites in ESCC and CTRL. **(i)** Distribution of RNA editing sites across genomic regions in ESCC and CTRL. **(j)** Categorization of variants in non-coding and coding regions in ESCC and CTRL or known and novel groups. **(k)** Sequence logo surrounding adenosine editing sites (±4 bp). The probability of nucleotides in the sequence logo showing frequency of each base for upstream (+4 bp) and downstream (−4 bp) of the editing sites.

Ultimately, a total of 222,020 high-confidence RNA editing sites were retained across ESCC and CTRL samples, including 124,486 mismatches unique to ESCC and 88,596 mismatches shared with CTRL (**Supplementary Table 2**). Principal component analysis (PCA) depicted a clear separation between ESCC and CTRL samples, indicating distinct RNA editing profiles associated with tumor status (**Fig. 1f**). Most of base substitutions occurred from adenosine to inosine, and the abundance of RNA mismatches in ESCC identified was two-fold than in CTRL (69% vs. 31%) (**Fig. 1g**). The quantity of edits varied across chromosomes (**Supplementary Fig. 2a**), and most genes had at least 2 edits per gene (**Supplementary Fig. 2b**). Notably, RNA editing frequency in 3′ UTR, non-coding RNAs (ncRNAs), and introns were significantly increased in ESCC compared to CTRL (**Supplementary Fig. 2c**), suggesting a dysregulated RNA editome in tumors. Up to 95% edits were categorized to known SNV edits in REDIportal [37], while 5% represented novel editing sites (**Fig. 1h**). Approximately 90% of editing sites resided within short interspersed nuclear element (SINE), yielding a total of 89,369 and 198,034 mismatch locations in CTRL and ESCC, respectively (**Fig. 1i**). Most of mutations were detected within intronic areas, followed by ncRNAs, 3′ UTRs and intergenic regions, with only a small fraction occurring in exons (**Fig. 1j**).

To examine the preference nucleotide surrounding RNA editing sites, sequence contexts surrounding edits were analyzed. Result revealed that nucleotide upstream adenosine (+1) was enriched in uridine whereas downstream position (−1) was depleted in guanosine (**Fig. 1k**), aligning with known ADAR substrate preferences [38]. Intriguingly, four consensus motifs ‘WGUAGAU’, ‘UGUAAUU’, ‘GWGUAGW’, and ‘AGUAGKW’ were strongly enriched in normal tissues, while only ‘GWGUAGW’ remained highly enriched in tumor samples (**Supplementary Fig. 3a, b**). These features suggested the reliability and accuracy of the mismatches identified using the combined tools, alongside the sequence context, which validated variants we detected.

### Widespread RNA editing reprogramming in primary ESCC with limited alterations during lymph node metastasis

To understand RNA editing in ESCC, we compared editing levels across ESCC and CTRL samples. We observed that RNA editing rates across genes and chromosomes were significantly increased in ESCC compared to CTRL (**Fig. 2a**). RNA editing levels for A-to-I substitutions depicted significant up-regulation across genes and patients (**Fig. 2b,c**). As it is known that higher coverage yields more RNA editing sites, we questioned if the rise of RNA editing level might result from increased coverage. Our data revealed that coverage of each site in ESCC remained unaltered and decreased in patients relative to CTRL (**Fig. 2b,c**). Besides, the editing rate of novel edits was significantly higher than known edits while read coverage remain unchanged (**Fig. 2d**).

**Fig. 2.**
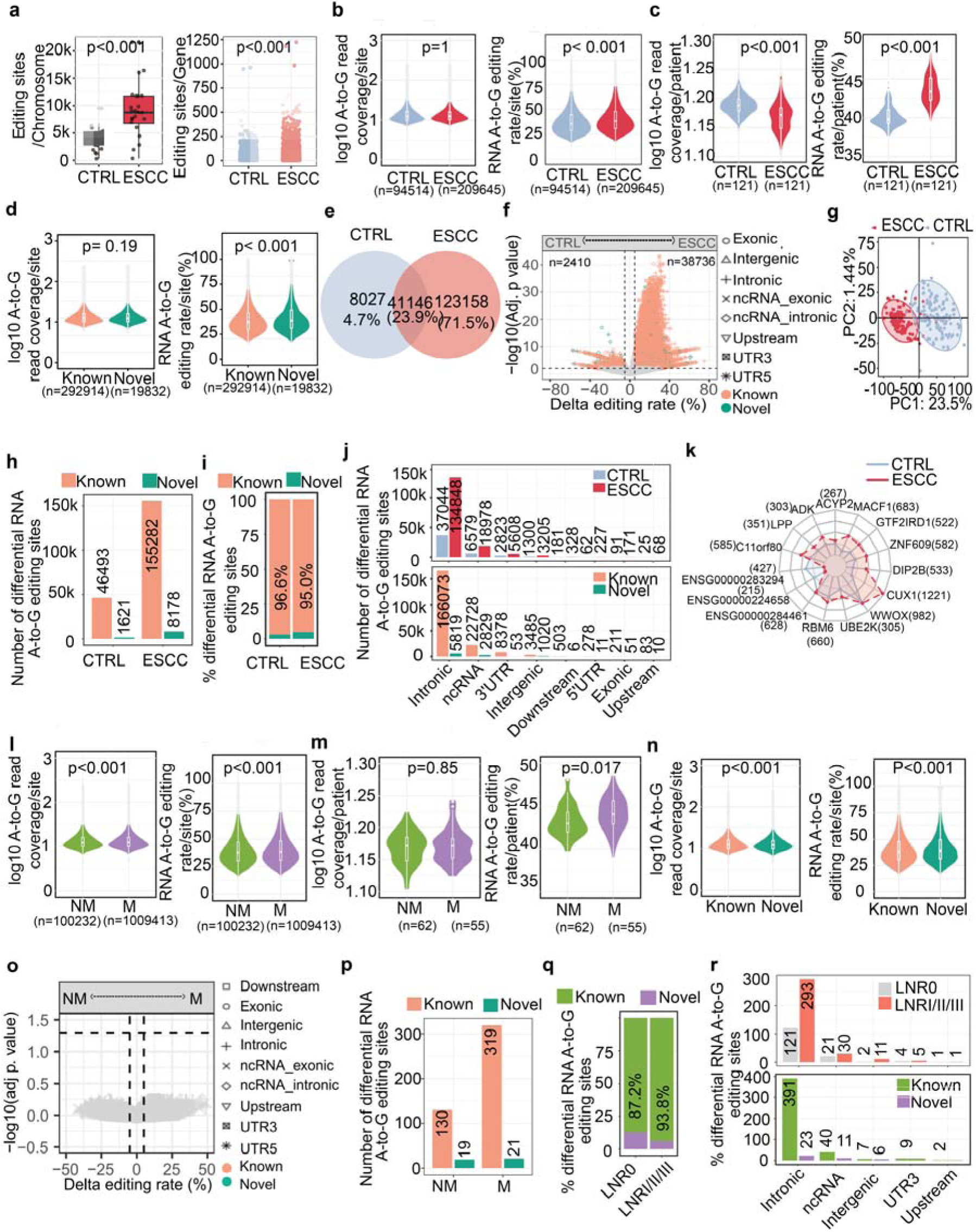
Genome-wide RNA editing landscape reveals enhanced editing rate in ESCC. **(a)** RNA editing sites in chromosomes or genes. **(b)** Read coverage for A-to-I edits or patients across ESCC and CTRL samples. **(c)** RNA A-to-I editing coverage or rate for patients cross ESCC and CTRL samples. **(d)** Read coverage or editing rate for A-to-I edits across known and novel groups. **(e)** Venn diagram showing RNA edits present in ESCC and CTRL samples. **(f)** Volcano plot presenting shared differentially edited sites (DRS) in ESCC and CTRL. Shapes represent distinct variant types in ESCC and CTRL samples. **(g)** PCA analysis showing differential A-to-I editing sites present in more than 50% samples. **(h,i)** The abundance of known and novel DRS A-to-I editing sites in ESCC and CTRL samples. **(j)** Distribution of variant types for DRS A-to-I edits in ESCC and CTRL samples, and DRS A-to-I edits that were known and novel. **(k)** Enrichment of genes containing DRS A-to-I edits in ESCC and CTRL samples. Genes containing the top 10 abundance of edits and corresponding editing counts were visualized. **(i,m)** Read coverage for A-to-I edits or patients in non-metastatic (NM) and metastatic (M) samples. **(n)** Read coverage or editing rate for A-to-I edit in known and novel groups. **(o)** Volcano plot presenting shared DRS in ESCC and CTRL samples. Shapes represent distinct variant types in NM and M samples. **(p)** The abundance of known and novel DRS A-to-I editing sites in NM and M samples. **(q)** Distribution of DRS A-to-I edits in NM and M samples, and DRS A-to-I edits that were known and novel. **(r)** Distribution of variant types for ESCC-and CTRL-specific A-to-I editing sites, classified as known or novel. Significant differences were indicated between groups. P values were calculated by the Wilcoxon test.

To identify differentially edited A-to-I sites, we performed pairwise comparisons and identified a total of 172,331 differentially RNA edited sites (DRSs). Among these, 71.5% (n = 123,158) were ESCC-specific editing events, whereas only 4.7% (n = 8,027) were CTRL-specific (**Fig. 2e; Supplementary Table 3**). In addition, 41,146 DRSs were shared between ESCC and CTRL. Of these shared sites, 2,410 exhibited significantly decreased editing rates in ESCC, whereas the vast majority (n = 38,736) showed significantly increased editing rates (**Fig. 2f)**. Notably, these DRSs alone were sufficient to robustly distinguish ESCC from CTRL samples, indicating a profoundly reprogrammed RNA A-to-I editing landscape in ESCC (**Fig. 2g)**. Furthermore, 163,565 DRSs corresponded to previously annotated editing sites, while 8,766 represented novel editing events (**Fig. 2h,i**). The preferential localization of DRSs within intronic regions **(Fig. 2j)** indicates that dysregulated RNA editing is more likely to influence gene expression via post-transcriptional regulation—especially alternative splicing and transcript maturation—rather than by directly reshaping protein-coding sequences.

It was shown that the RNA editing rates of most shared edits in ESCC cohort were above 30%, and an extraordinary number of edits were driven by ESCC (**Supplementary Fig. 2d,e**). 4,256 genes harboring differential A-to-I RNA editing sites in CTRL (n = 49,173 sites) and 6153 in ESCC (n = 164,304 sites) samples. Further analysis summarizes the top 15 genes with the highest burden of DRSs, revealing broadly increased RNA editing levels in these genes in ESCC relative to CTRL (**Fig. 2k**). We also examined the relationship between differential editing sites and *ADARs* expression. Of note, the number of differentially edited sites decreased significantly when *ADAR1* expression was included as a covariate (n = 5,370 sites) (**Supplementary Fig. 4a**). In contrast, *ADAR2* and *ADAR3* covariates influenced a small subset of differential editing sites (n = 21,285 and n = 22,707 sites, respectively) (**Supplementary Fig. 4b,c**), suggesting a dominant dependency of differential editing on *ADAR1* activity.

To determine whether RNA editing dysregulate during metastasis, ESCC were further stratified into two subgroups: non-metastatic and metastatic tissues. We found that RNA editing rates were enhanced during metastasis in sites (**Fig. 2l and Supplementary Fig. 5a)** and patients (**Fig. 2m)**, and dominantly located in introns (**Supplementary Fig. 5b**) with elevated coverage in metastatic ESCC compared to non-metastatic samples (**Fig. 2l**). Both coverage and editing rates were higher in novel edits (**Fig. 2n**). Despite the identification of non-shared DRSs between metastatic and non-metastatic cohorts (**Fig. 2o**), only several hundred cohort-specific RNA editing events were detected, with a modest enrichment in lymph node-metastatic cases (**Fig. 2p-r**). These observations indicate that aberrant RNA editing may exert a more substantial influence during primary ESCC tumorigenesis than during later metastatic progression.

Further result showed that RNA editing distribution is unlikely affected by gene length due to its minimal correlation with RNA editing distribution (R^2^ = 0.031 in CTRL and R^2^ = 0.043 in ESCC) (**Supplementary Fig. 6a**). Enrichment for gene analysis (**Supplementary Fig. 6b,c**) showed that 190 genes (out of 5104) enriched with DRS, in which 61 genes belong to DEGs (DEG1; **Supplementary Table 4**). To figure out whether genes harboring DRS correlate with differentially expressed genes, we conducted fisher’s exact test comparing two datasets. Result showed that up to 9.8% genes harboring DRS tended to be DEGs (p<0.0001, odds ratio=1.39; **Supplementary Fig. 7a**). Next, we examined the relationship between gene expression levels and RNA editing events. In both ESCC and CTRL samples, genes harboring at least one RNA editing site were predominantly concentrated in the mid-to-high expression bins (log2 normalized counts 8-14), with ESCC samples exhibiting a greater number of edited genes across almost all expression levels compared with CTRL (**Supplementary Fig. 7b**). Classification of RNA editing sites as known or novel revealed that most sites were annotated as known across all expression bins, whereas novel sites were relatively rare but consistently observed, particularly in the mid-expression bins (**Supplementary Fig. 7c**). These results indicate that RNA editing preferentially targets moderately to highly expressed genes, with novel events representing a smaller but detectable fraction, and is modestly associated with differential gene expression while being largely independent of gene length.

### Clinical relevance of RNA recoding A-to-I editing in primary and metastatic ESCC

To investigate DRSs that induce amino acid substitutions, we identified editing sites resulting in missense mutations. This analysis revealed 39 editing events affecting 25 genes, of which 11 genes harbored exclusively ESCC-specific sites, one gene contained only CTRL-specific sites, and 13 genes harbored sites shared between ESCC and CTRL (**Fig. 3a and Supplementary Table 5**). Then, we wonder whether there is any preference for amino acids within genes containing recoding sites. We found 39 recoding sites resulting in only 10 amino acid changes (**Fig. 3b**). Three substitutes, Gly (n = 11), Arg (n = 9), and Val (n = 6), accounted for up to 26 sites out of 39 (66.7%), suggesting site-selective editing towards mRNAs in ESCC and CTRL. RNA editing of these genes containing recoding sites displayed higher pattern in ESCC in comparison to control samples (**Fig. 3c**). Two of the occurrences of mismatches *c.A1984G* (I662V) in NBPF8 (novel edits), and *c.A308G* (Q103R) in CDK13 (known edits) with a high incidence rate in more than 80% patients, were validated by cloning and Sanger sequencing (**Fig. 3d**).

**Fig. 3.**
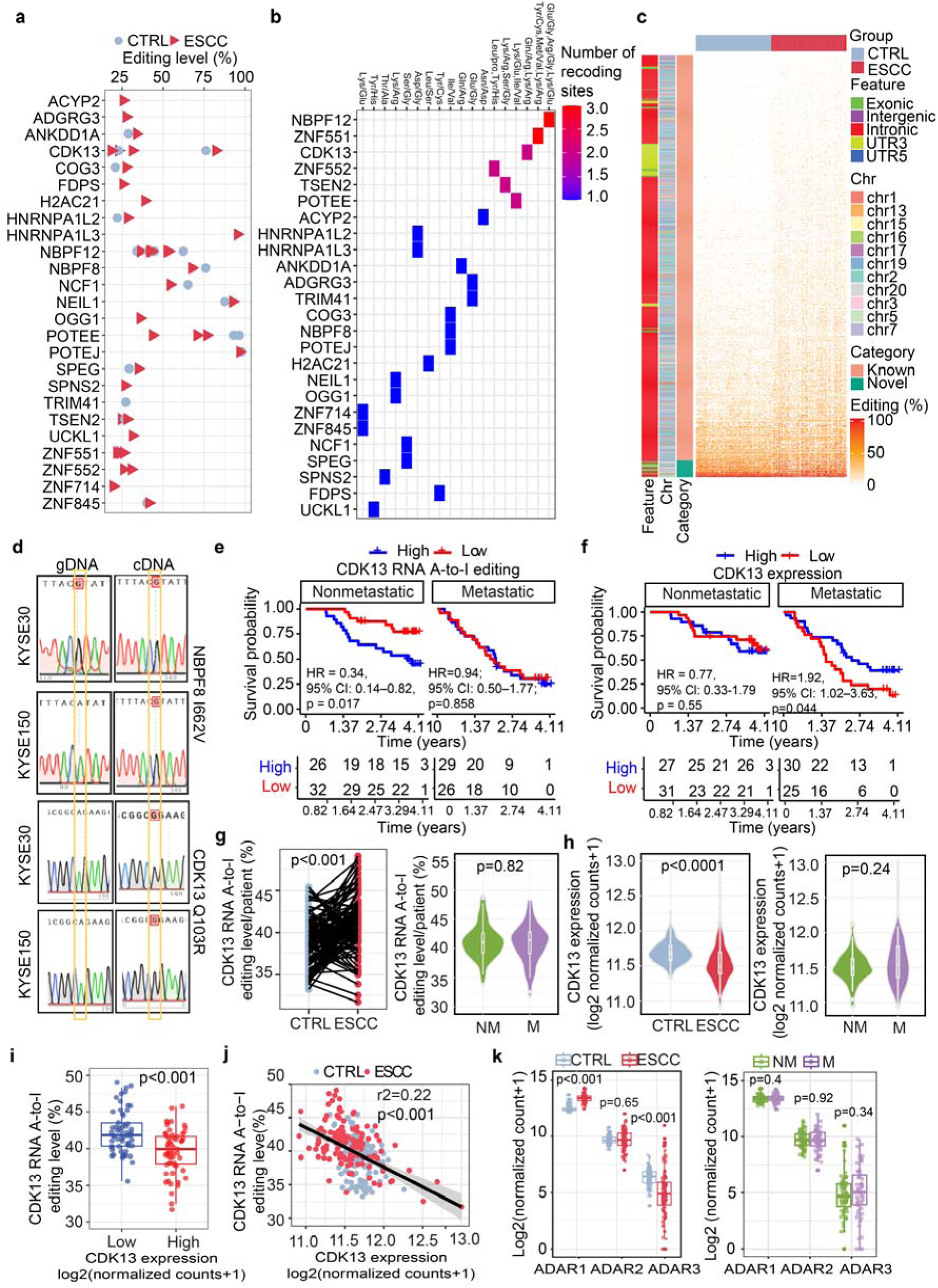
RNA editing profiles in recoding genes harboring differentially edited sites across ESCC and CTRL samples and their clinical relevance. **(a)** Dot plot showing editing levels of differentially edited recoding sites in genes in ESCC and CTRL. **(b)** Amino acid preferences of differentially edited recoding sites in genes. **(c)** Heatmap showing RNA editing levels of recoding genes containing at least one differentially edited recoding site. **(d)** Sanger sequencing chromatograms for two differentially edited recoding sites at position 1984 in *NBPF8* and position 308 in *CDK13*. The orange box highlights the peak of adenosine or guanosine. Edited nucleotide was shaded in red. **(e,f)** Kaplan-Meier curves for survival of high RNA A-to-I editing **(e)** and expression **(f)** versus low *CDK13* editing and expression cohorts in NM and M ESCC, respectively. NM: nonmetastatic, M: metastatic. P values were assessed by Cox regression. **(g)** Overall RNA A-to-I editing levels of *CDK13* across ESCC and paired CTRL samples, and across NM and M ESCC, respectively. (h) *CDK13* expression across ESCC and CTRL, and NM and M samples. **(i)** RNA A-to-I editing in *CDK13* in two groups of high and low *CDK13* expression based on the mean. **(j)** Correlation analysis of RNA A-to-I editing and expression of *CDK13*. **(k)** Expression of *ADARs* across samples of ESCC and CTRL. Significant differences were assessed by the Wilcoxon test.

To explore whether genes containing RNA editing sites that reprogram proteome associate with patient survival, we conducted Kaplan-Meier survival analysis linking overall RNA editing level to patient survival outcomes(**Supplementary Fig. 8a**). Stratification of ESCC patients by metastatic status revealed that single *CDK13* recoding sites showed no significant association with survival in either the non-metastatic or metastatic cohorts (**Supplementary Fig. 8b**). However, in the non-metastatic cohort, patients with high *CDK13* editing exhibited significantly worse survival compared to those with low editing (HR = 0.34, 95% CI 0.14-0.82, log-rank p = 0.017), whereas survival remained comparable between editing groups in the metastatic cohort (**Fig. 3e**). In contrast, survival of high editing in *OGG1* is associated with increased mortality risk during metastasis (**Supplementary Fig. 8a**). The remaining 23 genes with multiple recoding sites showed no significant survival differences, though some, such as *SPEG*, *COG3*, and *NBPF12*, displayed non-significant trends (**Supplementary Fig. 8a**).

Interestingly, high *CDK13* expression, while not associated with survival in non-metastatic patients, was significantly negatively correlated with poor prognosis in metastatic patients (**Fig. 3f**). Further analysis showed that *CDK13* editing levels were significantly elevated in ESCC tissues compared to CTRL, but did not differ between non-metastatic and metastatic ESCC (**Fig. 3g**), while *CDK13* expression was markedly reduced in tumors relative to CTRL, with no difference between non-metastatic and metastatic cases (**Fig. 3h**). These findings suggest that early ESCC development may be driven by high *CDK13* editing and concomitant down-regulation, whereas metastasis is more likely driven primarily by *CDK13* down-regulation. Consistently, *CDK13* editing levels were inversely correlated with its expression (**Fig. 3i,j**). Together, these results delineate a stage-dependent regulatory pattern in ESCC, in which hyper-editing of *CDK13* predominates during early tumor development, whereas disease progression in metastatic ESCC relies more heavily on *CDK13* downregulation.

### *CDK13* RNA editing is predominantly driven by *ADAR1* in ESCC

ADAR proteins are known active RNA A-to-I editing enzymes [39]. To investigate potential mediators of *CDK13* hyper-editing, we examined *ADARs* expression and found that *ADAR1* was up-regulated in ESCC, *ADAR3* was down-regulated, and *ADAR2* remained stable; none of the three changed during metastasis (**Fig. 3k**). Additionally, both *ADAR1* isoforms (P110 and P150) were up-regulated in tumors (**Supplementary Fig. S9a**). Further analyses suggest that RNA A-to-I editing levels and events were strongly and positively correlated with *ADAR1* expression, whereas no significant association was observed with *ADAR2*, and a negative correlation was detected with *ADAR3* (**Supplementary Fig. 9b,c**). Similarly, *ADAR1* expression was positively correlated with overall A-to-I editing and editing events in *CDK13*, whereas *ADAR3* showed a negative association, and no significant correlation was observed with *ADAR2* (**Fig. 4a**), and both *ADAR1* isoforms positively correlated with editing in *CDK13* (**Supplementary Fig. 9d**). Furthermore, we performed correlation analysis between the editing levels of all RNA editing sites and the expression of *ADAR1*, *ADAR2*, and *ADAR3* (**Supplementary Table 6**), and identified two recoding sites in *CDK13* (Q35R and Q103R) that were positively correlated with *ADAR1*expression (**Supplementary Fig. 9e**).

**Fig. 4.**
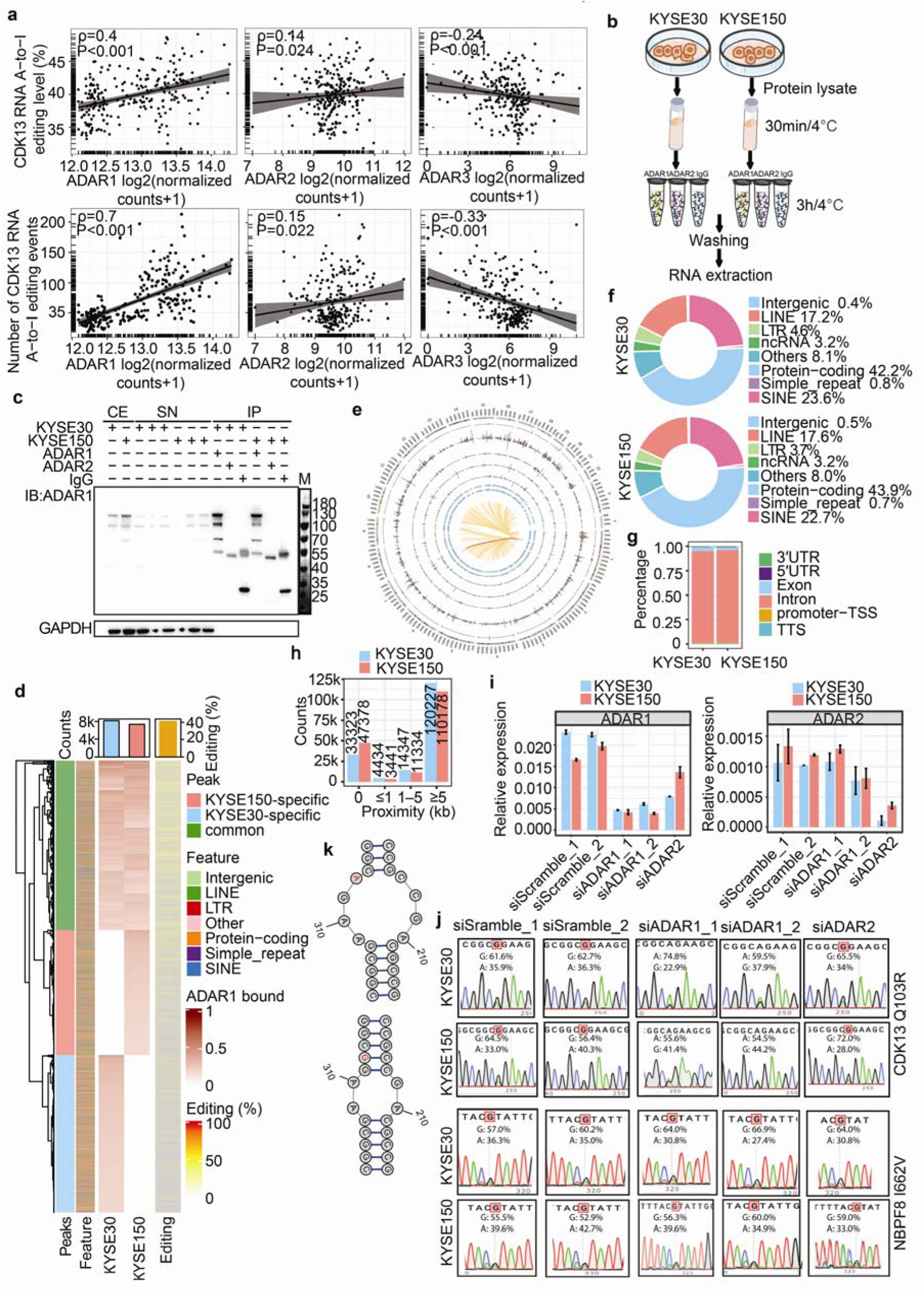
RNA editing in *CDK13* mainly depends on *ADAR1* activity. **(a)** Correlation of the RNA editing rates (upper panel) or RNA editing events (lower panel) in *CDK13* with the expression of *ADAR1, ADAR2,* and *ADAR3*. **(b)** Illustration of RNA immunoprecipitation (RIP) using KYSE30 and KYSE150 cells. Cells were collected and then equally split into three parts for agarose-conjugated antibody incubation, followed by washing and RNA extraction. **(c)** Immunoblotting examining ADAR1 abundance in distinct fractions of ADAR1 immunoprecipitation. GAPDH was used as loading control. Abbreviations represent CE: crude extract; SN: supernatant; IP: Immunoprecipitation. **(d)** Distribution of peaks (KYSE30 and KYSE150) and DRS across genomics elements. **(e)** Circos plot showing the hg38 genomic landscape of A-to-I editing events. From outermost to innermost, tracks 1-5 showed the editing densities (sites per million base pairs) in tumor (red bars) versus adjacent normal (blue bars) tissues for exons, introns, UTRs, ncRNAs, and intergenic regions, respectively; tracks 6-7 displayed scatter plots of peak scores for KYSE30 and KYSE150 cell lines. Peaks were annotated according to its genomic location, with red dots (exonic), blue (intronic), green (regulatory), and grey (intergenic). The inner links showed the genes correlated with ADAR in A-to-I editing, and the link to CDK13 was highlighted. **(f)** Distribution of RNA editing sites across genomics elements in two KYSE cells. **(g)** Distribution of RNA editing sites resided in different elements of protein-coding RNAs. **(h)** Counts of A-to-I DRS within or outside peaks. **(i)** Relative expression of *ADAR1* and *ADAR2* in siRNA *ADAR1*, *ADAR2* or control (Scramble) knockdown cells. Error bars represent ±S.D. of the mean from two technical replicates**. (j)** Sanger sequencing chromatograms for two recoding sites CDK13 Q103R (*CDK13 c.A308G*) and NBPF8 I662V (*NBPF8 c.A1984G*) were examined from cDNA of siRNA *ADAR1*, *ADAR2* or Scramble knockdown cells. **(k)** RNA secondary structure of unedited (upper panel) and edited (low panel) sequence surrounding *c.A308G* in *CDK13*.

To determine ADAR1 interacting RNAs, we performed RNA-immunoprecipitation (RIP) in two ESCC cell lines (KYSE30 and KYSE150) using antibodies against ADAR1, ADAR2 and IgG. The latter two antibodies served as controls, and the precipitated RNAs of ADAR1 were analyzed by RNA sequencing (**Fig. 4b,c and Supplementary Fig. 10**), whereas those of ADAR2 and IgG yielded too little RNAs for library construction. We identified 9,732 and 7,698 high signal peaks in KYSE30 and KYSE150, respectively (**Fig. 4d and Supplementary Fig. 11a**). The Circos plot depicting the landscape of A-to-I editing events in ESCC closely resembles the A-to-I editing pattern detected by *ADAR1* RIP in the two ESCC cell lines, highlighting the robustness of the analysis (**Fig. 4e**). Consistently, most peaks fell into introns at protein-coding RNAs and SINE (**Fig. 4f,g**). Approximately 18% (n = 33,323 sites in KYSE and n = 47,378 sites in KYSE150) of DRS located within peaks (**Fig. 4h**).

To discover conserved ADAR1 binding regions, 6,149 overlapping peaks from two cell lines were retained which were mapped in 3,712 genes (**Supplementary Table 7**). 39.2% (n = 2,412) peaks were classified to transposable element and 44.3% (n = 2,721) belong to protein-coding RNA (**Supplementary Fig. 11b**). Many DRS overlapped with these peaks across all genomes except for chromosome Y (**Supplementary Fig. 11c**). ADAR1 bound transcripts were subsequently identified, which participate in cell cycle related processes such as mitotic cytokinesis and chromatid segregation, and microtubule binding, indicating these genes may play crucial roles in cytoskeletal dynamics and chromosome segregation (**Supplementary Fig. 11d**).

*CDK13* contains a total of 288 RNA A-to-I edits, including 284 intronic mismatches and 4 exotic mismatches. Only 133 intronic variants located within peaks and the rest of them (121 intronic edits and 4 recoding sites) were beyond peaks. To examine experimentally whether direct interacting at editing sites with ADAR1 is required, PCR amplification and Sanger sequencing surrounding recoding sites in *CDK13* and *NBPF8* (control) were conducted after inhibiting *ADAR1* and *ADAR2* expression in *siADAR1* or *siADAR2*-treated lines (**Fig. 4i**). Result showed that knockdown of *ADAR1* eliminated editing at the *CDK13 c.A308G* (Q103R) site (**Fig. 4j**) in the distal stem loop of binding regions that located outside of peaks in two cells (**Fig. 4k**). Unedited CDK13 transcripts had an MFE of −203.70 kcal/mol and ensemble diversity of 70.95, while editing decreased the MFE to −218.10 kcal/mol and increased ensemble diversity to 106.32 (**Fig. 4k**), indicating that A-to-I editing may modulate local RNA structure and protein binding, though its effect on overall transcript stability requires further experimental confirmation.

### *ADAR1*-mediated *CDK13* editing forms a feedback loop promoting ISG signaling in early ESCC

Given that a large number of RNA editing sites, including two high-frequency recoding events, were detected in *CDK13* in more than 80% of ESCC patients, and that elevated *CDK13* editing levels were significantly associated with poor survival during ESCC progression, we hypothesized that A-to-I editing of *CDK13* pre-mRNA may regulate its function at the post-transcriptional level, thereby contributing to ESCC tumorigenesis. Consistent with this hypothesis, our analyses demonstrated a significant negative correlation between *CDK13* A-to-I editing levels and *CDK13* expression (**Fig. 3i,j**). Moreover, comparative analyses between tumor tissues and matched adjacent normal tissues showed that *ADAR1* expression was markedly upregulated in ESCC (**Supplementary Fig. 9a**), whereas *CDK13* expression was significantly downregulated (**Fig. 3h**). To further investigate the crosstalk between *ADAR1* and *CDK13*, we knocked down CDK13 in ESCC cell lines and examined the expression of ADAR family genes (**Fig. 5a,b**). *CDK13* depletion specifically increased *ADAR1* expression, without affecting *ADAR2* (*ADARB1*) or *ADAR3* (*ADARB2*), suggesting that loss of *CDK13* function releases a suppressive constraint on *ADAR1* (**Fig. 5a,b**). These results collectively suggest that *ADAR1*-mediated A-to-I editing of *CDK13* forms a regulatory crosstalk, in which reduced *CDK13* expression relieves suppression of ADAR1, potentially contributing to ESCC tumorigenesis.

**Fig. 5.**
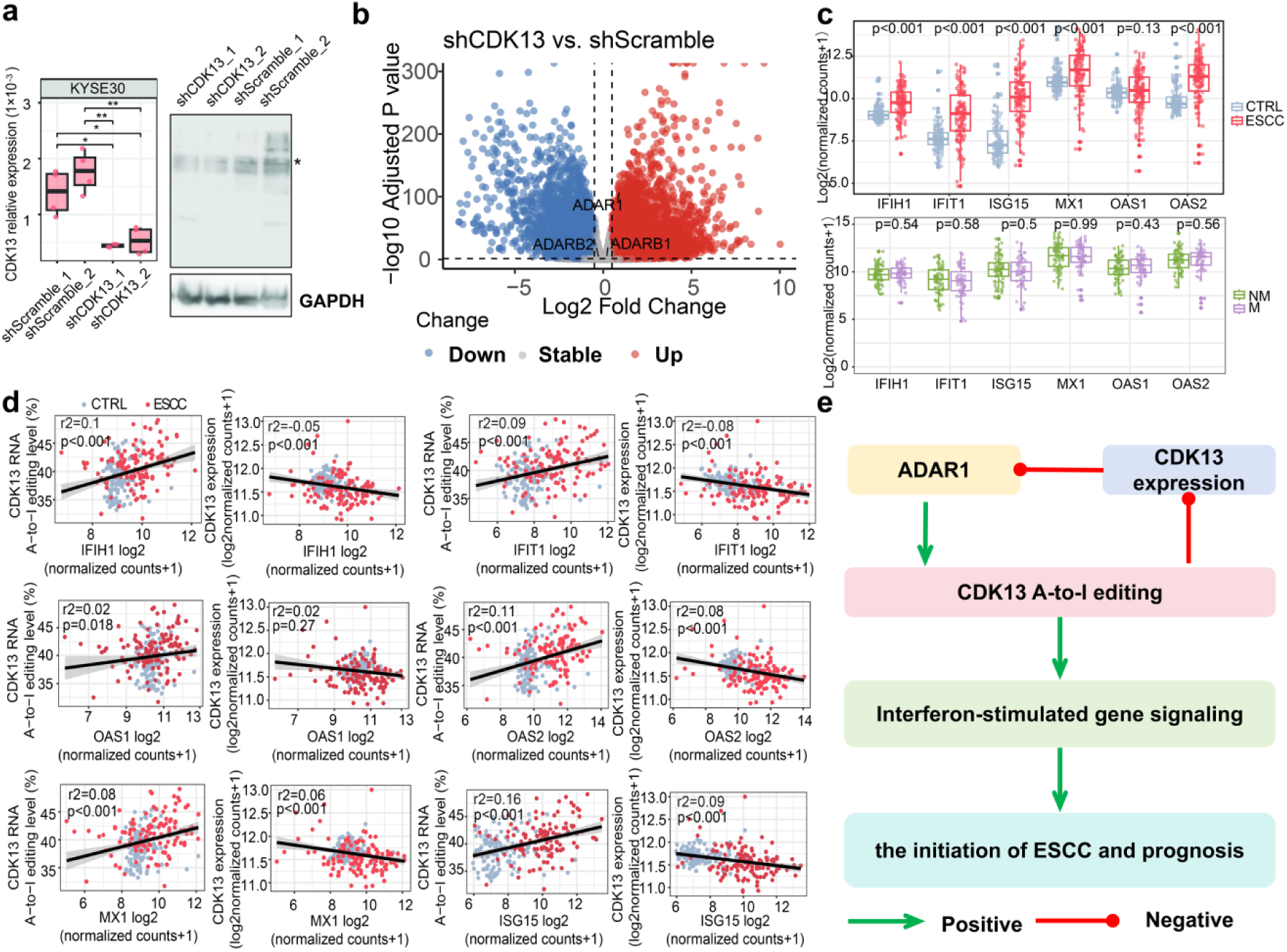
*CDK13* knockdown and its impact on *ADAR1* associated with interferon-stimulated genes (ISGs) in ESCC. **(a)** Relative *CDK13* expression in two stable shRNA knockdown and control KYSE30 cell lines (left) and Western blot of CDK13 in the same cell lines (right); asterisks indicate CDK13. **(b)** Volcano plot showing differential gene expression upon *CDK13* knockdown in KYSE30 cells, with ADAR family members highlighted. **(c)** Expression of representative ISGs in ESCC versus control (top) and in primary versus metastatic tumors (bottom). **(d)** Regression analysis of six ISGs with *CDK13* RNA A-to-I editing and *CDK13* expression. **(e)** Schematic illustration of ADAR1/CDK13 regulation in ESCC prognosis. Data are mean ± S.D.; significance was assessed by Student’s t-test. Asterisks denote significance (*p < 0.05, **p < 0.01, ***p < 0.001).

Genetic or pharmacological inactivation of *CDK12* and its paralog *CDK13* has been shown to robustly induce cGAS–STING-mediated antitumor immunity in preclinical models [40]. In our study, we observed a distinct molecular state in ESCC characterized by elevated *CDK13* RNA editing accompanied by reduced *CDK13* expression (**Fig. 3g-j**). This prompted us to investigate the expression of key interferon-stimulated genes (ISGs), which serve as readouts of cGAS – STING signaling. Unexpectedly, six key ISGs were significantly upregulated in ESCC relative to matched controls (**Fig. 5c**). Notably, their expression showed no significant association with lymph node metastasis status (**Fig. 5c**), consistent with our previous observation that *CDK13* editing levels were not correlated with lymph node metastasis. Consistent with the above observation, correlation analyses showed that *CDK13* editing levels were positively associated with ISG expression, whereas ISG expression was inversely correlated with *CDK13* expression (**Fig. 5d**). Together, these findings indicate that *ADAR1*-mediated A-to-I editing of *CDK13* establishes a positive feedback loop that modulates ISG signaling (**Fig. 5e**), potentially sustaining a chronic, submaximal activation of the cGAS – STING pathway and contributing to immune evasion and ESCC progression.

### Multi-omics dissects an *ADAR1-CDK13* phospho-signaling axis linking cytoskeletal remodeling, trafficking, and cGAS–STING signaling in ESCC

To investigate whether RNA A-to-I editing of *CDK13* contributes to ISG signaling regulation, we performed integrated transcriptomic and phosphoproteomic analyses in stable *CDK13*-knockdown KYSE30 cells (**Supplementary Fig. 12a,b**). BigWig analysis revealed a marked reduction in read density upon *CDK13* knockdown (**Fig. 6a**). Phosphoproteomic profiling identified over 3,000 phosphorylated proteins (**Supplementary Fig. 12c**), together with 302 differentially expressed genes (DEG2; **Fig. 6b and Supplementary Table 8**) and 1,638 differential phosphorylation sites (DPS), comprising 847 hyperphosphorylated and 791 hypophosphorylated sites (**Fig. 6c and Supplementary Table 9**).

**Fig. 6.**
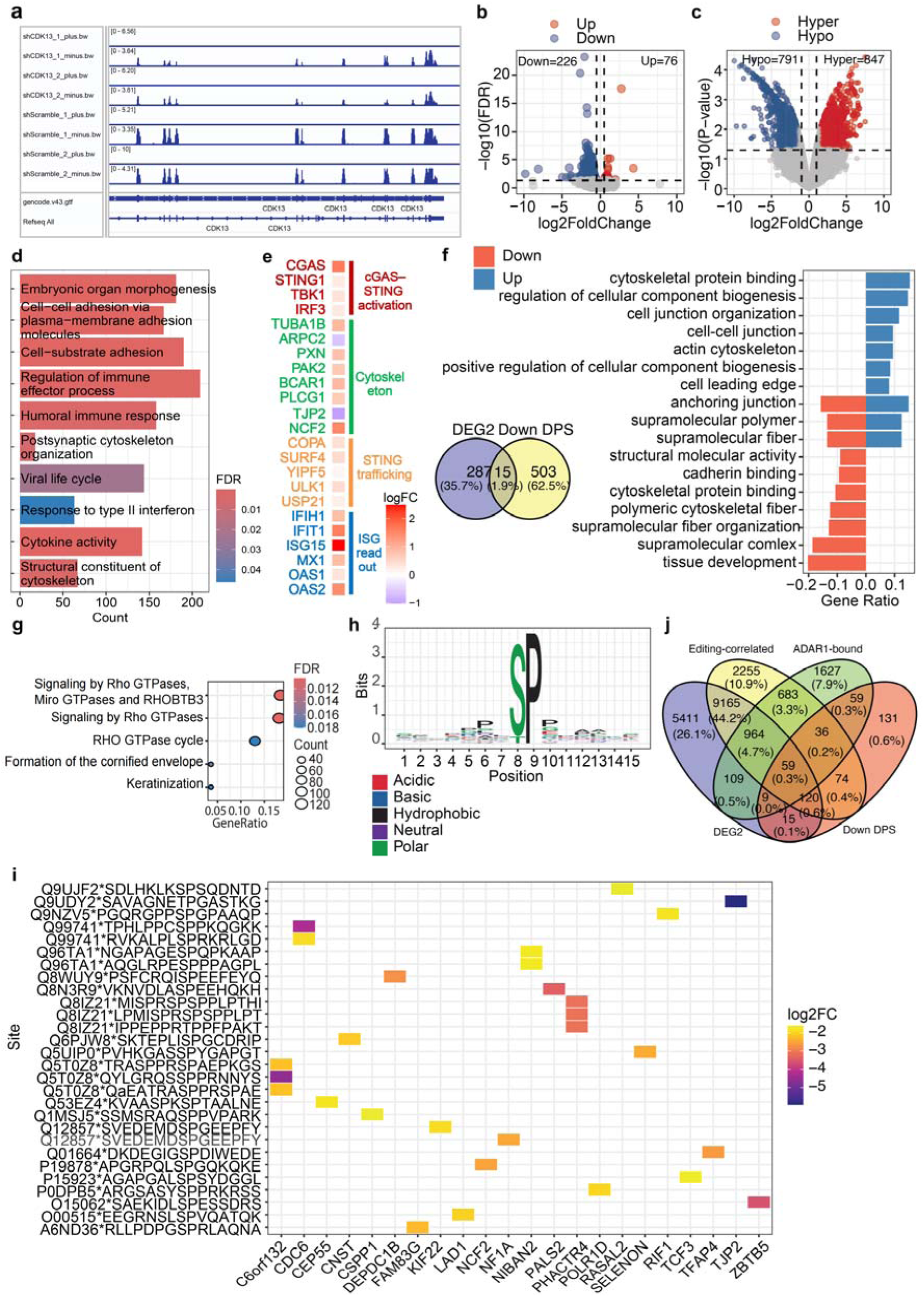
Integrative analysis of CDK13 knockdown reveals RNA A-to-I editing-dependent regulation of phosphorylation, cytoskeleton, and STING-mediated innate immune signaling in ESCC. **(a)** Reduced *CDK13* signal in stable shRNA knockdown KYSE30 cells compared with controls. rRNA-depleted reads were separated by strand and normalized to CPM. **(b)** Volcano plot of differentially expressed genes (DEG2) in *CDK13* knockdown versus control cells. DEG2 were defined as log2FC > 0.5 or < −0.5 with FDR < 0.05. Blue and red dots indicate down- and up-regulated DEG2, respectively. **(c)** Volcano plot of differentially phosphorylated sites (DPS). DPS were defined as log2FC > 1.5 or < −1.5 with p < 0.05. Blue and red dots represent hypo- and hyper-phosphorylated sites, respectively. **(d)** DEG2 is significantly enriched in immune response, cytoskeleton, and interferon signaling pathways. **(e)** Heatmap showing fold changes (logFC) of genes related to cGAS – STING activation, cytoskeleton regulation, STING trafficking, and interferon-stimulated gene (ISG) readout upon *CDK13* knockdown in KYSE30 cells. **(f)** Venn diagram of DEG2 and hypo-phosphorylated sites, and the top 10 GO terms enriched among non-DEG2 DPS ranked by FDR. FDR was controlled at 5%. Blue and orange bars indicate GO terms enriched in proteins containing up- and down-phosphorylated sites, respectively. **(g)** Dot plot of Reactome pathways enriched for down-regulated DPS. **(h)** Potential CDK13 targets involved in ADAR1-dependent RNA editing. Twenty-two proteins containing 28 DPS were identified with the canonical proline residue directly following the phosphorylated amino acid**. (i)** Sequence conservation surrounding phosphorylated residues (±7 amino acids). Colors indicate amino acid chemistry, and bits represent residue conservation. **(j)** Venn diagram integrating four datasets: ADAR1-binding genes (from overlapping peaks), RNA editing-correlated genes (DEG1 and editing-correlated), DEG2, and down-regulated DPS.

Functional enrichment analysis showed that the DEGs were predominantly associated with immune-related pathways, cell adhesion, and cytoskeleton organization (**Fig. 6d and Supplementary Fig. 12d,e**). Consistently, *CDK13* knockdown resulted in robust upregulation of core components of the cGAS–STING pathway, including *CGAS* and *STING*, as well as enhanced expression of downstream ISG readouts (**Fig. 6e and Supplementary Fig. 12e**). In parallel, genes involved in cytoskeletal remodeling and STING trafficking were also significantly upregulated (**Fig. 6e**). Although STING is classically a cytosolic DNA sensor triggering type I interferon and NF-κB signaling, its activation increasingly appears to depend on cytoskeletal dynamics and intracellular trafficking, which in turn regulate cytokine expression and cell adhesion [41]. Together, these data indicate that *CDK13* knockdown is associated with activation of the cGAS–STING signaling axis, potentially through cytoskeletal reorganization and facilitated STING trafficking.

To delineate the post-translational mechanisms underlying this process, we focused on proteins exhibiting downregulated phosphorylation sites while maintaining unchanged RNA expression levels (n = 503; **Fig. 6f**). Enrichment analyses revealed that these proteins were significantly involved in cytoskeleton organization, cell adhesion and anchoring junctions, and Rho GTPase-related signaling pathways (**Fig. 6f,g**). To further discover how these proteins were regulated by editing in CDK13 pre-mRNAs, we linked 503 proteins (harboring 791 down-regulated phosphorylation sites) with differentially expressed genes (**Supplementary Table 4**), editing-correlated genes (**Supplementary Table 10**), and ADAR1-bound RNAs (**Supplementary Table 7**). This analysis identified 59 proteins that were both closely associated with RNA editing and bound by ADAR1 in ESCC compared with controls (**Fig. 6j**). Among these, nine proteins—including TUBA1B, ARPC2, PAK2, TJP2, NCF2, PLCG1, PXN, MAK13, and BCAR1—were enriched in IgSF cell adhesion molecule signaling (CAM) or leukocyte transendothelial migration (**Supplementary Fig. 13**).

Given that CDK family kinases preferentially phosphorylate Ser/Thr residues followed by a proline (S/T-P motif) [42], we further incorporated motif preference and evolutionary conservation criteria, identifying 28 phosphopeptides from 22 proteins as putative CDK13 substrates (**Fig. 6h,i and Supplementary Fig. 12f**). Notably, these 22 proteins were consistently enriched in pathways related to cytoskeleton organization, cell adhesion, and anchoring junctions (**Supplementary Fig. 12g**), supporting a role for *CDK13* in modulating structural and trafficking-related processes. Together, these findings suggest that RNA editing-dependent regulation of CDK13 substrates orchestrates cytoskeletal remodeling, trafficking, and cGAS–STING signaling in ESCC.

## Discussion

RNA editing diversifies the transcriptome and proteome, and it is globally present in animals, plants, insects and the sexual stage of fungi. The most prevalent base alteration in mammals is A-to-I changes, which dominantly depends on enzymatic activity of ADARs. Although many A-to-I conversions occur in repetitive regions and introns, some mutations reside in exons, leading to missense alterations and involvement in the central neural system [43–45]. However, few studies characterize global analysis of RNA editing in tumors and matched normal tissues and provide overview for potential mRNA candidates of ADARs. Whether RNA editing plays a role in ESCC carcinogenesis and metastasis remains unclear. We wonder how the dysregulated RNA editome influences humans with ESCC or other diseases.

In this study we conducted whole-transcriptomic sequencing in ESCC and paired control cases from 121 individuals to analyze the RNA editome. We have identified a total of 222,020 high-confidence RNA SNVs from ESCC and CTRL tissues and found that RNA editing level was elevated in localized and metastatic ESCC compared to paired normal tissues, showing heterogeneous pattern of editing sites across samples. Our study revealed the global landscape of RNA editing in ESCC and the DRS reprograming protein diversity, with 124,486 ESCC-sepcific sites. However, although a massive number of sites were differentially edited in early-stage ESCC, only condition-specific sites were identified as differentially edited during metastasis. This suggested the pivotal role of RNA A-to-I editing in the localized ESCC.

We have identified many DRS by pairwise comparison in ESCC and control, but a limited amount when comparing NM and M samples. 12 mutations are predicted to be pathogenic according to ProtVar (**Supplementary Table 5**). However, whether these recoding sites are pathogenic or benign requires further experimental verification. Evidence showed CDK13 interacted with splicing factor ASF/SF2 associated p32 [46] and L-type cyclins [47], regulating alternative splicing nearly two decades ago. CDK13 is known to be essential in more than eleven diseases [48–52]. It cooperates with CDK12 to regulate transcription via RNA polymerase II [53] and is required for ZC3H14 phosphorylation for nuclear RNA surveillance. CDK13 involves in nuclear RNA homeostasis which regulates the accumulation of aberrant [54]. Knockdown of *CDK13* attenuates cell viability and accelerates melanoma in zebrafish [54]. Recently, *CDK13* was reported to regulate lipid metabolism [55] and to phosphorylate 4E-BP1 [56]. Inhibitors targeting CDK12/CDK13 have been developed for the treatment of breast cancer [49] and ovarian cancer [57]. Surprisingly, we found editing in *CDK13* among 55 RNA missense mutations was uniquely associated with the prognosis of primary ESCC, indicating that RNA editing is likely play vital roles in primary ESCC but is less important in metastasis. ADAR1 driven RNA editing abundance of *CDK13* may serve as a promising biomarker for early prognosis in localized ESCC, whereas decreased expression of *CDK13* may promote tumor progression in advanced stages of the disease. Whether complement edited *CDK13* influences cell proliferation and migration is still unknown and further exploration needs to be conducted. Therefore, *CDK13* functions as a context-dependent regulator in cancer, whose dysregulation may contribute to tumor initiation or progression through perturbation of transcriptional and post-transcriptional homeostasis.

Considering CLIP-seq introduces bias for certain RNA substrates and increases non-specific binding and may somehow damage cellular status [58], there native RIP-seq was exploited coupling with RNA editing uncovering ADAR1 catalyzed editing sites. We confirmed knockdown of ADAR1 decrease editing efficiency in KYSE30 and KYSE150 cells. Recently several RNA edits resided in exons of *CDK13* were reported [59, 60], but how does ADAR1 edited *CDK13* influence cellular processes in tumors needs to be explored which may be exploited for therapeutic avenue. Here we characterized candidates catalyzed by ADAR1 using overlapping peaks of two ESCC cells mapped to genes with RNA editing data. 5 genes harboring differentially edited sites causing missense mutation were identified present in peaks calling from RIP-seq of KYSE30 and KYSE150, including *CDK13*, *HNRNPA1L2*, *NBPF12*, *NCF1* and *ZNF714* potentially as conservative and high fidelity ADAR1 substrates.

Although previous studies reported that acute inactivation of *CDK12/13* elicits robust STING-mediated antitumor immunity in preclinical models [40], our findings in human ESCC reveal that ADAR1-mediated CDK13 editing forms a feedback loop promoting ISG signaling in early ESCC (**Fig. 5c-e**). Importantly, *CDK13* knockdown in ESCC cells similarly induced cGAS – STING and ISG signaling, potentially through cytoskeletal reorganization and enhanced STING trafficking, supporting a cell–intrinsic response (**Fig. 6d,e** and **Supplementary Fig. 12d,e**). We propose that RNA editing-mediated partial attenuation of *CDK13* establishes a chronic, non-lethal ISG program that promotes immune evasion and tumor initiation, rather than effective antitumor immunity. Consistently, previous work [61] has shown that loss of *ADAR1* in tumor cells reduces A-to-I editing of interferon-inducible RNAs, enhancing dsRNA sensing and cell–intrinsic inflammatory signaling, which underscores the role of RNA editing in restraining innate immune activation and supports our model in which *ADAR1*-mediated *CDK13* editing establishes chronic ISG activation. Our study contributes to elucidating the physiological and pathological meaning of RNA editing in the axis of *ADAR1* with *CDK13* in early tumorigenesis.

Leveraging RNA-seq data from 121 ESCC and control samples, phosphorproteomics and RIP-seq, 59 proteins were identified that were significantly associated with RNA editing, differentially expressed in DEG1, contained down-regulated phosphorylation sites and were bound by ADAR1 (**Fig. 6j**). Among these, nine proteins —TUBA1B, ARPC2, PAK2, TJP2, NCF2, PLCG1, PXN, MAK13, and BCAR1— were enriched in IgSF CAM signaling and leukocyte transendothelial migration pathways, highlighting a broader involvement of cytoskeletal and adhesion remodeling in *CDK13*-associated immune regulation.

The global RNA editing burden of *CDK13* closely correlated with prognosis in primary ESCC, which probably through regulating phosphorylation of 22 proteins (**Fig. 6i**), which contain canonical motif for proline-directed kinases with a residue proline at downstream of phosphorylation site S/T (**Fig. 6h**). Their phosphorylation levels were down-regulated in *CDK13* knockdown cells with unchanged transcript abundance. These findings suggest that they are likely regulated by CDK13 and play meaningful role in RNA editing. Whether they are directly phosphorylated by CDK13 as non-canonical substrates other than RNA polymerase II [53] and translational machinery component 4E-BP1 [56] requiring further experimental exploration. In addition, 10 out of 22 proteins encompassing RASAL2, TJP2, NFIA, PALS1, SELENON, PHACTR4, KIF22, CSPP1, NIBAN2, LAD1 may play roles in maintaining tissue structure and organization of supramolecular assemblies.

Our study provides a comprehensive and integrative view of the RNA A-to-I editing landscape in ESCC and reveals a previously unappreciated role of the *ADAR1-CDK13* axis in shaping tumor-intrinsic immune signaling. By combining transcriptomics, phosphoproteomics, and RIP-seq analyses, we demonstrate that ADAR1-driven RNA editing selectively targets *CDK13*, leading to its functional attenuation and rewiring of downstream phosphorylation networks involved in cytoskeletal organization, RNA homeostasis, and innate immune regulation. Distinct from acute CDK12/13 inactivation that elicits robust antitumor immunity [40], RNA editing–mediated partial suppression of *CDK13* induces persistent, cell – intrinsic cGAS–STING activation, potentially promoting immune evasion and early tumor initiation in ESCC. These findings position *CDK13* not as a canonical oncogene, but as a context-dependent regulatory hub whose dysregulation via RNA editing perturbs transcriptional and post-transcriptional homeostasis. Together, our work highlights RNA editing as a critical and dynamic layer of gene regulation in ESCC pathogenesis and suggests that editing-dependent vulnerabilities within the *ADAR1-CDK13* axis may represent promising avenues for biomarker development and therapeutic intervention in early-stage disease.

## Methods

### Sample collection, RNA library preparation and sequencing

ESCC and matched peritumoral control (CTRL) tissues were obtained from 127 patients who had not undergone radiotherapy or chemotherapy before surgery at the First Affiliated Hospital of Zhengzhou University. The study was approved by the Ethics Committee of Zhengzhou University [62]. Tumor samples were further classified as localized (n = 62) or lymph node-positive stage I-III disease (LI, n = 23; LII, n = 18; LIII, n = 14) based on the number of involved lymph nodes. High-quality total RNA was extracted using TRIzol Reagent (Invitrogen) according to the manufacturer’s protocol. RNA-seq libraries were prepared using the VAHTS® Total RNA-seq (H/M/R) Library Prep Kit for Illumina® and sequenced on an Illumina X Ten platform (BGI) to generate 2 × 150 bp paired-end reads with strand-specific (reverse) orientation. Raw sequencing data were subjected to quality assessment using FastQC (v0.11.8) [63]. Samples with excessively low read counts were excluded, including two paired ESCC-adjacent normal samples and four CTRL samples lacking matched ESCC counterparts. After quality control and sample filtering, a total of 121 ESCC tumors with matched normal tissues were retained for all subsequent analyses.

### Cell culture

Human ESCC cell lines KYSE30 (CL-0577) and KYSE150 (CL-0638) were purchased from Procell Co. Ltd (Wuhan, China). The cells were cultured in RPMI 1640 medium (Gibco, C11875500BT) supplemented with 10% fetal bovine serum (FBS) (Zeta Life, Z7181FBS-500) in a 37°C incubator with 5% CO_2_. When collecting the cells, they were treated with trypsin-EDTA for 2-3 min, and the reaction was stopped by 2 volumes of medium supplemented with 10% FBS. The cells were then suspended in FBS containing 10% DMSO (Sigma-Aldrich) and stored in a cell freezing program at −80°C for two days and were transferred to liquid nitrogen for long-term storage.

### RT-qPCR

Total RNA was extracted using the RNA Easy Fast Cell Kit (TIANGEN, DP451-TA), followed by DNase I treatment (Invitrogen, 18068015). cDNA synthesis was performed using Superscript III Reverse Transcriptase (Invitrogen, 18080093) and oligo(dT) primer. qPCR was conducted using TB Green® Premix Ex Taq TM II (Takara) and amplified in a CFX96™ System (Rio-Rad). Relative transcript levels were calculated using the 2^-ΔCt^ and normalized by housekeeping genes (*HRPT1*/*ACTB*/*TBP*). Results were representative of three independent experiments. Primer oligos were synthesized by GENEWIZ and are listed in **Supplementary Table 11**.

### Genomic DNA extraction

Genomic DNA was extracted using 2% CTAB as previously described [64]. Briefly, homogenized cells were incubated in a 60°C incubator for 40 min prior to chloroform: isoamyl alcohol (24:1) extraction and Isopropanol precipitation in a −20°C freezer for 1 h. The DNA pellet was washed with 75% ethanol and then dissolved in 190 µL DEPC-treated water. 10 µL RNase A (10 mg mL^-1^, Tiangen, Y2221) was added to remove RNA contaminants and the reaction was incubated at 70°C for 15 min, followed by overnight ethanol precipitation in a −20°C freezer. On the other day, DNA pellet was collected by centrifugation and was washed with 75% ethanol, air-dried, and finally dissolved in DEPC-treated water.

### Unsupervised analysis

The regularized logarithm transformation (rlog)-transformed read counts were applied to principal component analysis (PCA). Multidimensional scaling (MDS) analysis was performed based on the rlog-transformed data to assess the sample similarity and clustering. RNA editing sites that were missing or undetected in more than 50% (n = 60) of samples were removed from the PCA analysis to reduce noise and improve interpretability.

### Pre-processing, RNA editing detection, variants filtering and gene-based annotation

Quality trimming of all raw FASTQ files were performed using Trim-Galore (v0.5.0) and reads were aligned to the GENCODE human genome assembly (GRCh38.p13) [65] using STAR (v2.7.10b) [66] with 2-pass mapping which allows splice awareness, followed by rRNA removal. Gene or transcript count matrices for ESCC and CTRL samples were generated using RSEM (v1.3.1) [67]. 5′ end (6-nt) reads were clipped to exclude hexamer primers using trimBam from bamUtil toolkit (v1.0.15) [68] and reverse-strand reads were extracted for further processing. PCR duplicates introduced during library preparation were removed using Picard tools (v2.27.5). To further increase the calling accuracy before variant detection, aligned reads were preprocessed using the GATK toolkits (v3.8 and v4.3) [69], including ReorderSam, SortSam, AddOrReplaceReadGroups, MarkDuplicates, SplitNCigarReads, IndelRealigner, BaseRecalibrator, and ApplyBQSR. Finally, variant detection was performed using bcftools (v1.17) [70] mpileup and call.

ANNOVAR (v2022-08-02) [71] was leveraged to retrieve genomic features and annotated variants that lead to amino acid substitutions. Recoding variants were further verified by CADD (v1.7) [72] and examined in ProtVar (v1.4) [73] to check whether they are pathogenic or benign. To obtain RNA-DNA mismatches with high confidence, we applied several stringent filtering criteria to the variants. We required that (i) the variants have a minimum read coverage of 5, a minimum calling score of 30, and a minimum supporting read count of 3; (ii) the variants with a single alternative base at the same site and an allele frequency of at least 0.1; (iii) variants that overlapped with genomic SNPs from 1000 Genome Project [74], dbSNP (db138) [75] and the University of Washington Exome Sequencing Project (ESP) [76] were excluded; iv) multiallelic sites and variants located within 4 bp of splice junctions were discarded; and v) candidate variants must present in more than 5% of the samples (121, n = 6). Known RNA editing sites were identified by intersecting with SNPs from REDIportal [77]. Editing sites absent in the REDIportal were labeled as novel sites.

### Quantification of RNA editing levels, RNA editing events and read coverage

RNA editing levels were defined as the percentage of alternative reads in the total number of aligned reads for each genomic site and patient using the following formula: RNA editing level = ∑edited reads / ∑total reads. RNA editing events were defined as the total number of non-missing edits for each patient. Read coverage was calculated as the total number of aligned reads spanning each site: Read coverage = ∑total reads.

### Identification of differential RNA editing sites

Differentially editing sites between ESCC and CTRL samples were identified using a linear model with donor ID as a covariate (limma (v3.54.2) [78]). The expression of *ADAR1*, *ADAR2*, and *ADAR3* was covaried distinctively to measure their influence on RNA editing patterns. All significance values were adjusted for multiple testing using the Benjamin-Hochberg (BH) method to control the false discovery rate (FDR). Sites passing delta(Δ) RNA editing rate > 5 or < −5 and a FDR < 0.01 were labeled significant.

### Analysis of RNA editing by gene length

A pairwise comparison across ESCC and CTRL groups was conducted to analyze differential A-to-I RNA editing. The number of RNA editing events per gene was normalized by the log-transformed gene length (number of edits per gene/log2(gene length +1)) to remove the potential effects of gene length on editing frequency. Outliers were identified as genes with normalized RNA editing counts falling outside the 99% confidence intervals (CIs) of the mean. Enriched genes were defined as outliers based on values exceeding the 99% confidence threshold.

### Gene Ontology term and Reactome pathway enrichment

Gene Ontology (GO) and Reactome pathway enrichment analysis were performed using the R packages clusterProfiler (v4.6.2) [79] and ReactomePA (v4.6.2) [80]. The Benjamin-Hochberg method was applied to control the FDR with a significance threshold of 5%. Expressed genes detected in RNA-seq or phosphoproteomics were used as background sets.

### Differentially expressed genes identification

The R package DESeq2 (v1.40.2)[81] was used to transform the raw read counts for differentially expressed genes (DEGs). For RNA seq data of 121 ESCC and paired CTRL samples, genes with log2 fold change (log_2_FC) > 0.5 or < −0.5 and FDR < 0.01 were considered as DEGs (DEG1). For mRNA seq data of shRNA targeting *CDK13* and control Scramble cell lines, genes with log_2_FC > 0.5 or < −0.5 and FDR < 0.05 were defined as DEGs (DEG2).

### Two-sided Fisher’s exact test

To explore the correlation between differentially edited sites (DRS) and differentially expressed genes (DEG1), we conducted two-sided Fisher’s exact test. The number of genes detected in RNA seq of ESCC and matched peritumor tissues (ESCC vs. CTRL) was used as background set.

### Native RNA immunoprecipitation sample preparation

Harvested cells were lysed and disrupted in 12 mL IP extraction buffer (20 mM Tris-HCl, 300 mM NaCl, 5 mM MgCl_2_, 0.5% (v/v) NP40, 5 mM DTT, Protease inhibitor (cOmplete, Roche), Ribolock RNase inhibitor (Thermo Scientific)) for 30 min on a rotating wheel. Protein lysates were collected by centrifugation at 12,000 × g for 10 min and the supernatant was transferred to a 5 mL tube to remove cell debris. 200 μL crude extract of KYSE30 and KYSE150 was taken for WB. 12 mL of crude extract was split evenly to three parts in 5 mL tube to incubate with 30 µg pre-cleared ADAR1 AC (Santa Cruz, sc-73408 AC) and ADAR2 AC (Santa Cruz, sc-73409 AC) antibodies as well as normal mouse IgG AC (Santa Cruz, sc-2343 AC) as a negative control. Protein lysates were then incubated on a roller wheel for 3 hours at 4°C in a refrigerator. After centrifugation at 300 × g for 30 s, 200 μL IP supernatant were taken for WB. The rest of the wash supernatant was removed and agarose beads were re-suspended in 1 mL IP washing buffer (20 mM Tris-HCl, 300 mM NaCl, 5 mM MgCl_2_, 0.5% (v/v) Triton X-100, 5 mM DTT, Protease inhibitor, Ribolock RNAse inhibitor). The captured complexes were transferred to new 1.5 mL tubes and washed 5 times gently and collected by centrifugation at a speed of 200 × g, 4°C for 30 s. In the final step, washing buffer was removed and agarose beads were re-suspended in 1 mL washing buffer. The beads were then split to two portions including 300 μL washing buffer containing ADAR1-conjugated agarose for WB as the IP fraction, and 700 μL for RNA isolation.

### RNA isolation from RNA immunoprecipitation

RNA was extracted from agarose-conjugated (AC) ADAR1 or normal mouse IgG AC. Agarose gels with ADAR1-RNA complex were re-suspended in 150 µL RNA release buffer (100 mM Tris-HCl (pH 7.5), 10 mM EDTA, 300 mM NaCl, 2% SDS, 1 µg/µL proteinase K). RNA was released from antibody bound agarose beads by incubation at 60°C and 300 rpm thermomixer for 15 min prior to isolation with citrate-saturated phenol. Centrifugation was performed at the speed of 10,000 × g at 4°C for 10 min, and the upper aqueous layer was transferred to a new 1.5 mL Eppendorf (EP) tube and mixed with phenol/methylene chloride/Isoamyl alcohol mixture (25:24:1), followed by 10 times inversion and centrifugation at 4°C, 10,000 × g at 4°C for 10 min. The upper aqueous was transferred to a new 1.5 mL EP tube and mixed with methylene chloride/Isoamyl alcohol mixture (24:1) by 10 times inversion. This step was repeated twice and the upper aqueous was transferred to a new 1.5 mL EP tube and precipitated with ethanol and 20 g Glycogen RNA grade in a −20°C freezer overnight until further usage.

### Western blotting

For RNA immunoprecipitation, agarose beads were re-suspended in 1 × SDS loading buffer (P0015F, Beyotime) diluted with washing buffer, heated at 100°C for 5 min, and immediately cooled on ice for 10 min. For stable knockdown cell samples, protein extraction was performed as described in the section phosphor-proteomic analysis. Proteins were resolved on a Bis-Tris SDS-PAGE gel (GenScript, M00654) using electrophoresis buffer (GenScript, M00138) at 100 volts for 1 h 40 min at 4°C. Subsequently, proteins were transferred to a methanol-activated PVDF membrane at 50 mA overnight (16 h) at 4°C. The membrane was blocked with 5% non-fat milk powder in 1 × TBST (Biocomma, P164230701) for 1-2 hours at room temperature (RT), followed by incubation with primary anti-ADAR1 antibody (Santa Cruz, sc-73408; 1:200 dilution) overnight (∼16 h) in 4°C refrigerator. After three times washes with 1 × TBST (10 min), the membrane was incubated with secondary anti-mouse antibody (Abcam, ab205719; 1:2000 dilution) in 5% non-fat milk powder in 1 × TBST for 1-2 h at RT. Following three washes with 1 × TBST (10 min), protein signals were detected using enhanced chemiluminescence (ECL, MeilunBio, MA0186). PVDF membrane was then stripped using stripping buffer (Thermo Scientific, 46430) and reprobed with primary antibody against GAPDH (Proteintech, 60004-1-Ig; 1:5000 dilution) following standard protocols.

For transient knockdown cells, harvested cells were lysed in RIPA buffer (Beyotime, P0013C) with Protease Inhibitor. Protein concentrations were quantified using the BCA assay (Thermo Scientific, A65453). ∼20 µg of proteins per sample were loaded onto a PAGE gel (EpiZyme, PG112), and resolved using electrophoresis buffer (Beyotime, P0014D) at 100 V for 1 h 40 min at 4°C. Proteins were then transferred to a methanol-activated PVDF at 300 mA for 3 h at 4°C. The membrane was blocked with 5% non-fat milk powder in 1 × TBST for 1-2 hours at RT, followed by overnight incubation (∼16 h) at 4°C refrigerator with primary antibody against ADAR1 (Santa Cruz, sc-73408; 1:200 dilution) or CDK13 (Bethyl, A301-458A-BL; 1:2000 dilution). After three washes with 1 × TBST, the membrane was incubated with secondary antibody against mouse (Abcam, ab205719; 1:2000 dilution) or rabbit (Proteintech, SA00001-19; 1:2000 dilution) with 5% non-fat milk powder in 1 × TBST for 1-2 hours at RT. protein signals were detected using enhanced chemiluminescence. PVDF membrane was then stripped using stripping buffer (Thermo Scientific, 46430) and reprobed with primary antibody against GAPDH (Proteintech, 60004-1-Ig; 1:5000 dilution) following standard protocols.

### RNA immunoprecipitation library construction, Illumina sequencing, reads quality control and mapping

The fragment distribution, intactness and concentration of purified RNAs from agarose beads were assessed using an Agilent 2100 Bioanalyzer (Agilent) and a simpliNano spectrophotometer (GE Healthcare). Immunoprecipitated RNAs with good quality were used for subsequent library preparation by Novogene Co. Ltd (Beijing, China) using NEB Next^®^ Ultra™ RNA library Prep Kit (New England Biolabs). Library quality was examined on the Agilent 2100 system. Qualified libraries were sequenced on an Illumina NovaSeq X Plus platform with 2 × 150 bp unstranded paired-end reads according to the standard protocols. After Trim-Galore trimming to remove adapters and low-quality reads from fastq files, clean reads were aligned to the GENCODE human genome (GRCh38.p13) using STAR in 2-pass mode. Reads not paired properly, and with mapping quality bellow 20 were filtered. rRNA were removed before calling peaks using MACS2 (v 2.2.7.1) [82] with an FDR of 1%. Peaks related genes were confirmed by HOMER (v5.0) [83].

### RNA structure prediction

RNA secondary structure was predicted by RNAfold from ViennaRNA package [84]. The structure was visualized using StructureEditor 1.0 [85].

### Validation of RNA editing sites by PCR/cloning and Sanger sequencing

Putative RNA editing sites in *NBPF8* and *CDK13* were validated by cloning using genomic DNA and cDNA as the template, followed by Sanger sequencing. For editing efficiency in siRNA *ADAR1/2* knockdown cells, PCR amplicons amplified from cDNA were performed and subsequently subjected to Sanger sequencing. Primers used in PCR are listed in the **Supplementary Table 11**.

### Short interference RNA transfection

siRNAs targeting *ADAR1* (*siADAR1*_1, *siADAR1*_2), *ADAR2* (*siADAR2*), and *CDK13* (*siCDK13*_1, *siCDK13*_2), along with two non-targeting controls (*siScramble*_1, *siScramble*_2) (5 nM) were synthesized by TSINGKE Co. Ltd (Beijing, China). KYSE30 and KYSE150 cells were seeded at a density of 2 × 10^5^ cells per 60 mm petri dish in RPMI medium supplemented with 10% FBS and cultured at a 37°C incubator with 5% CO_2_ for 48 h. Cells were then transfected with oligonucleotides at a final concentration of 0.4 µM using lipofectamine RNAiMAX (Invitrogen, 13778150) according to manufacturer’s instructions. After 32-48 h of transfection, total RNA was extracted for downstream analysis, such as RT-qPCR or PCR amplification for Sanger sequencing. The sequences of siRNAs for *Scramble*_1 and *Scramble*_2, *ADAR1*_1, *ADAR1*_2, *ADAR2*, *CDK13*_1, *CDK13*_2, are listed following, respectively: (5′-UCUCUCACAACGGGCAU(dT)(dT), 5′-GUGGGCACCGAUAUCUUGA(dT)(dT), 5′-GACUAUCUCUUCAAUGUGU(dT)(dT), 5′-CUAUGAAAGCCAUGACAAU(dT)(dT), 5′-GAUCGUGGCCUUGCAUUAA(dT)(dT)) [86], 5′-GCCAGUGCAUCACAAACAA(dT)(dT), 5′-GCUGAAUUGAACAAGAAUA(dT)(dT)) [87].

### Lentiviral-genomic shRNA knockdown cells

Lentiviral particles carrying shRNA targeting *CDK13* and non-targeting control Scramble (4 ×10^8^ TU mL^-1^) were purchased from TSINGKE Co. Ltd. The virus was produced using the pLVX-U6-Puro transfer vector along with psPAX2 packaging and pMD2.G envelope plasmids. KYSE30 cells were transduced with lentivirus at a multiplicity of infection (MOI) of 80 in the presence of 8 μg mL^-1^ polybrene. After 48 hours, transduced cells were selected several rounds using 2-8 μg mL^-1^ puromycin for two weeks. Knockdown efficiency was validated by RT-qPCR and immune blotting.

### mRNA sequencing, reads quality control and mapping

Total RNA from cells with stable knockdown of *CDK13* (shCDK13) and control (shScramble) were isolated using RNA Easy Fast Cell Kit (TIANGEN, DP451-TA) with DNase I (TIANGEN, RT411) treatment according to the manufacturer’s instructions. Isolated total RNAs were examined by Agilent 2100 bioanalyzer (Agilent Technologies, A, USA) to assess RNA integrity and quantity. mRNA libraries were constructed and sequenced by Novogene Co. Ltd using Fast RNA seq lib prep kit v2 (ABclonal, rk20306). Libraries were examined using Qubit, RT-qPCR and Bioanalyzer. Qualified libraries were sequenced on an Illumina NovaSeq X Plus platform with 2 × 150 bp paired-end reads in a reverse strand-specific manner. Raw reads in fastq format were first processed using Trim-Galore to remove adapters and low-quality reads. Cleaned reads were then aligned to the GENCODE human genome (GRCh38.p13) using STAR in 2-pass mode, followed by the removal of unmapped reads and those with a mapping quality below 20. rRNA removed reads were then quantified using RSEM for downstream analysis.

### Quantitative phosphoproteomic analysis

Total protein was extracted from stable knockdown cells of *CDK13* and Scramble using IP extraction buffer (20 mM Tris-HCl, 300 mM NaCl, 5 mM MgCl_2_, 0.5% (v/v) NP40, 5 mM DTT, Protease inhibitor (Roche, 118361530), Ribolock RNAse inhibitor (Thermo Scientific, EO0382), PhosSTOP^TM^ (Roche, 4906845001)). Isolated proteins were split for immunoblotting and liquid chromatography-tandem mass spectrometry (LC-MS/MS). Digestion of proteins and enrichment of phosphorylated peptides were performed and analyzed by Novogene Co., Ltd using Vanquish^TM^ Neo UHPLC system coupled to an Orbitrap Astral^TM^ mass spectrometer (Thermo Scientific) in data-independent acquisition (DIA) mode. Peptide separation wa s performed on a C18 column (PrepMap^TM^ Neo, 150 µm × 15 cm, 2 µm particle size). Raw data were processed for peptide matching, site annotation, and quantification using Spectronaut. The resulting data were used for downstream analysis.

### Identification of differentially phosphorylated sites

Log-transformed phosphorylated sites were normalized to median, followed by fitting to a linear model for identification of differentially phosphorylated sites (DPS). Sites with log_2_FC > 1.5 or < −1.5 and p-value < 0.05 were defined as DPS.

### Identification of RNA editing correlated RNAs

Differentially expressed genes (DEG1) and genes contained RNA editing sites which were correlated to RNA editing were identified based on FDR < 0.05 and correlation coefficient (ρ) > 0.3 or < −3.

## Supplementary information

**Supplementary Figure 1-13**; **Supplementary Table 1.** Metadata of 121 paired samples used in this study; **Supplementary Table 2.** Table shows high confidence RNA-DNA mismatches identified in ESCC and paired adjacent normal tissues; **Supplementary Table 3**. Table shows high confidence differential RNA editing sites identified in ESCC and paired adjacent normal tissues; **Supplementary Table 4**. Table shows differential expression analysis of genes between ESCC and paired adjacent normal tissues; **Supplementary Table 5**. Table shows identified differential RNA A-to-I editing sites leading to missense mutation; **Supplementary Table 6**. Table shows correlation analysis of RNA editing level for identified RNA A-to-I editing sites with expression of ADARs; **Supplementary Table 7**. Table shows overlapping peaks in RIP-seq data of KYSE30 and KYSE150; **Supplementary Table 8**. Table shows differential expression analysis of genes between shCDK13 and paired control shScramble cell lines; **Supplementary Table 9**. Table shows differential phosphorylation analysis of sites between shCDK13 and control shScramble cell lines; **Supplementary Table 10**. Table shows correlation analysis of gene expression with average RNA editing level across samples; **Supplementary Table 11**. DNA and RNA Oligos used in the study.

## Authors’ contributions

W.Z. And L.H. conceived and designed the study. L.H. performed data curation, methodology development, formal analysis, validation, and visualization. LH and W.Z. drafted the manuscript. M.Y. and G.J. contributed to data curation, resources, validation, and manuscript revision. D.L. participated in data visualization and validation. W.Z. and G.J. supervised the study. W.Z., L.H., and M.Y. acquired funding. All authors reviewed and approved the final manuscript.

## Data availability

The whole-transcriptome sequencing data of ESCC tumors and CTRL were generated and deposited in the National Genomics Data Centre (NGDC), China (https://bigd.big.ac.cn/). The BioProject accession number is PRJCA001577, and the raw RNA-seq data are available in the Genome Sequence Archive (GSA) under the accession number HRA000111. Additional RNA sequencing data and quantitative phosphoproteomics data generated in this study have been deposited in the NCBI Sequence Read Archive (SRA) and GEO under BioProject PRJNA1398866, respectively.

## Declarations

### Ethics approval and consent to participate

This study was approved by the Ethics Committee of the First Affiliated Hospital of Zhengzhou University. Written informed consent was obtained from all participants at the time of enrollment. Clinical trial number: not applicable.

### Consent for publication

All authors contributed substantially to the conception, design, execution, and interpretation of the study, and reviewed and approved the final manuscript for publication.

## References

1. Bray F, Ferlay J, Soerjomataram I, Siegel RL, Torre LA, Jemal A. Global cancer statistics 2018: GLOBOCAN estimates of incidence and mortality worldwide for 36 cancers in 185 countries. CA Cancer J Clin 2018; 68(6):394–424.

2. Arnold M, Soerjomataram I, Ferlay J, Forman D. Global incidence of oesophageal cancer by histological subtype in 2012. Gut 2015; 64(3):381–7.

3. Jarmoskaite I, Li JB. Multifaceted roles of RNA editing enzyme ADAR1 in innate immunity. RNA 2024; 30(5):500–11.

4. Wang Q, Miyakoda M, Yang W, Khillan J, Stachura DL, Weiss MJ, et al. Stress-induced apoptosis associated with null mutation of ADAR1 RNA editing deaminase gene. J Biol Chem 2004; 279(6):4952–61.

5. Wang Q, Khillan J, Gadue P, Nishikura K. Requirement of the RNA editing deaminase ADAR1 gene for embryonic erythropoiesis. Science 2000; 290(5497):1765–8.

6. Tonkin LA, Saccomanno L, Morse DP, Brodigan T, Krause M, Bass BL. RNA editing by ADARs is important for normal behavior in Caenorhabditis elegans. EMBO J 2002; 21(22):6025–35.

7. Liddicoat BJ, Piskol R, Chalk AM, Ramaswami G, Higuchi M, Hartner JC, et al. RNA editing by ADAR1 prevents MDA5 sensing of endogenous dsRNA as nonself. Science 2015; 349(6252):1115–20.

8. Yuan J, Xu L, Bao HJ, Wang JL, Zhao Y, Chen S. Biological roles of A-to-I editing: implications in innate immunity, cell death, and cancer immunotherapy. J Exp Clin Cancer Res 2023; 42(1):149.

9. Burns CM, Chu H, Rueter SM, Hutchinson LK, Canton H, Sanders-Bush E, et al. Regulation of serotonin-2C receptor G-protein coupling by RNA editing. Nature 1997; 387(6630):303-8.

10. Berg KA, Cropper JD, Niswender CM, Sanders-Bush E, Emeson RB, Clarke WP. RNA-editing of the 5-HT(2C) receptor alters agonist-receptor-effector coupling specificity. Br J Pharmacol 2001; 134(2):386–92.

11. Price RD, Weiner DM, Chang MS, Sanders-Bush E. RNA editing of the human serotonin 5-HT2C receptor alters receptor-mediated activation of G13 protein. J Biol Chem 2001; 276(48):44663–8.

12. Visiers I, Hassan SA, Weinstein H. Differences in conformational properties of the second intracellular loop (IL2) in 5HT(2C) receptors modified by RNA editing can account for G protein coupling efficiency. Protein Eng 2001; 14(6):409–14.

13. Warhaftig G, Sokolik CM, Khermesh K, Lichtenstein Y, Barak M, Bareli T, et al. RNA editing of the 5-HT2C receptor in the central nucleus of the amygdala is involved in resilience behavior. Transl Psychiatry 2021; 11(1):137.

14. Yang JH, Sklar P, Axel R, Maniatis T. Editing of glutamate receptor subunit B pre-mRNA in vitro by site-specific deamination of adenosine. Nature 1995; 374(6517):77–81.

15. Rueter SM, Burns CM, Coode SA, Mookherjee P, Emeson RB. Glutamate receptor RNA editing in vitro by enzymatic conversion of adenosine to inosine. Science 1995; 267(5203):1491–4.

16. Melcher T, Maas S, Higuchi M, Keller W, Seeburg PH. Editing of alpha-amino-3-hydroxy-5-methylisoxazole-4-propionic acid receptor GluR-B pre-mRNA in vitro reveals site-selective adenosine to inosine conversion. J Biol Chem 1995; 270(15):8566–70.

17. Higuchi M, Single FN, Kohler M, Sommer B, Sprengel R, Seeburg PH. RNA editing of AMPA receptor subunit GluR-B: a base-paired intron-exon structure determines position and efficiency. Cell 1993; 75(7):1361–70.

18. Herb A, Higuchi M, Sprengel R, Seeburg PH. Q/R site editing in kainate receptor GluR5 and GluR6 pre-mRNAs requires distant intronic sequences. Proc Natl Acad Sci U S A 1996; 93(5):1875–80.

19. Davies B, Brown LA, Cais O, Watson J, Clayton AJ, Chang VT, et al. A point mutation in the ion conduction pore of AMPA receptor GRIA3 causes dramatically perturbed sleep patterns as well as intellectual disability. Hum Mol Genet 2017; 26(20):3869–82.

20. Kwak S, Kawahara Y. Deficient RNA editing of GluR2 and neuronal death in amyotropic lateral sclerosis. J Mol Med (Berl) 2005; 83(2):110–20.

21. Krampfl K, Schlesinger F, Zorner A, Kappler M, Dengler R, Bufler J. Control of kinetic properties of GluR2 flop AMPA-type channels: impact of R/G nuclear editing. Eur J Neurosci 2002; 15(1):51–62.

22. Greger IH, Khatri L, Ziff EB. RNA editing at arg607 controls AMPA receptor exit from the endoplasmic reticulum. Neuron 2002; 34(5):759–72.

23. Takuma H, Kwak S, Yoshizawa T, Kanazawa I. Reduction of GluR2 RNA editing, a molecular change that increases calcium influx through AMPA receptors, selective in the spinal ventral gray of patients with amyotrophic lateral sclerosis. Ann Neurol 1999; 46(6):806–15.

24. Chen L, Li Y, Lin CH, Chan TH, Chow RK, Song Y, et al. Recoding RNA editing of AZIN1 predisposes to hepatocellular carcinoma. Nat Med 2013; 19(2):209–16.

25. Shimokawa T, Rahman MF, Tostar U, Sonkoly E, Stahle M, Pivarcsi A, et al. RNA editing of the GLI1 transcription factor modulates the output of Hedgehog signaling. RNA Biol 2013; 10(2):321–33.

26. Lazzari E, Mondala PK, Santos ND, Miller AC, Pineda G, Jiang Q, et al. Alu-dependent RNA editing of GLI1 promotes malignant regeneration in multiple myeloma. Nat Commun 2017; 8(1):1922.

27. Tonkin LA, Bass BL. Mutations in RNAi rescue aberrant chemotaxis of ADAR mutants. Science 2003; 302(5651):1725.

28. Kawahara Y, Zinshteyn B, Sethupathy P, Iizasa H, Hatzigeorgiou AG, Nishikura K. Redirection of silencing targets by adenosine-to-inosine editing of miRNAs. Science 2007; 315(5815):1137–40.

29. Zhang L, Yang CS, Varelas X, Monti S. Altered RNA editing in 3’ UTR perturbs microRNA-mediated regulation of oncogenes and tumor-suppressors. Sci Rep 2016; 6(23226.

30. Qin YR, Qiao JJ, Chan TH, Zhu YH, Li FF, Liu H, et al. Adenosine-to-inosine RNA editing mediated by ADARs in esophageal squamous cell carcinoma. Cancer Res 2014; 74(3):840–51.

31. Fu L, Qin YR, Ming XY, Zuo XB, Diao YW, Zhang LY, et al. RNA editing of SLC22A3 drives early tumor invasion and metastasis in familial esophageal cancer. Proc Natl Acad Sci U S A 2017; 114(23):E4631–E40.

32. Hu Q, Chen Y, Zhou Q, Deng S, Hou W, Yi Y, et al. ADAR promotes USP38 auto-deubiquitylation and stabilization in an RNA editing-independent manner in esophageal squamous cell carcinoma. J Biol Chem 2024; 300(10):107789.

33. Ramaswami G, Zhang R, Piskol R, Keegan LP, Deng P, O’Connell MA, et al. Identifying RNA editing sites using RNA sequencing data alone. Nat Methods 2013; 10(2):128–32.

34. Bahn JH, Lee JH, Li G, Greer C, Peng G, Xiao X. Accurate identification of A-to-I RNA editing in human by transcriptome sequencing. Genome Res 2012; 22(1):142–50.

35. Levanon EY, Eisenberg E, Yelin R, Nemzer S, Hallegger M, Shemesh R, et al. Systematic identification of abundant A-to-I editing sites in the human transcriptome. Nat Biotechnol 2004; 22(8):1001–5.

36. Abdelwahab O, Belzile F, Torkamaneh D. Performance analysis of conventional and AI-based variant callers using short and long reads. BMC Bioinformatics 2023; 24(1):472.

37. Mansi L, Tangaro MA, Lo Giudice C, Flati T, Kopel E, Schaffer AA, et al. REDIportal: millions of novel A-to-I RNA editing events from thousands of RNAseq experiments. Nucleic Acids Res 2021; 49(D1):D1012–D9.

38. Bazak L, Haviv A, Barak M, Jacob-Hirsch J, Deng P, Zhang R, et al. A-to-I RNA editing occurs at over a hundred million genomic sites, located in a majority of human genes. Genome Res 2014; 24(3):365–76.

39. Ashley CN, Broni E, Miller WA. ADAR Family Proteins: A Structural Review. Curr Issues Mol Biol 2024; 46(5):3919–45.

40. Bao Y, Chang Y, Tien JC, Cruz G, Yang F, Mannan R, et al. CDK12/13 inactivation triggers STING-mediated antitumor immunity in preclinical models. J Clin Invest 2025; 135(18).

41. Taguchi T, Mukai K, Takaya E, Shindo R. STING Operation at the ER/Golgi Interface. Front Immunol 2021; 12(646304.

42. Moses AM, Heriche JK, Durbin R. Clustering of phosphorylation site recognition motifs can be exploited to predict the targets of cyclin-dependent kinase. Genome Biol 2007; 8(2):R23.

43. Chan TW, Fu T, Bahn JH, Jun HI, Lee JH, Quinones-Valdez G, et al. RNA editing in cancer impacts mRNA abundance in immune response pathways. Genome Biol 2020; 21(1):268.

44. Birgaoanu M, Sachse M, Gatsiou A. RNA Editing Therapeutics: Advances, Challenges and Perspectives on Combating Heart Disease. Cardiovasc Drugs Ther 2023; 37(2):401–11.

45. Li JB, Walkley CR. Leveraging genetics to understand ADAR1-mediated RNA editing in health and disease. Nat Rev Genet 2025; 26(8):532–46.

46. Even Y, Durieux S, Escande ML, Lozano JC, Peaucellier G, Weil D, et al. CDC2L5, a Cdk-like kinase with RS domain, interacts with the ASF/SF2-associated protein p32 and affects splicing in vivo. J Cell Biochem 2006; 99(3):890–904.

47. Chen HH, Wong YH, Geneviere AM, Fann MJ. CDK13/CDC2L5 interacts with L-type cyclins and regulates alternative splicing. Biochem Biophys Res Commun 2007; 354(3):735–40.

48. Sifrim A, Hitz MP, Wilsdon A, Breckpot J, Turki SH, Thienpont B, et al. Distinct genetic architectures for syndromic and nonsyndromic congenital heart defects identified by exome sequencing. Nat Genet 2016; 48(9):1060–5.

49. Quereda V, Bayle S, Vena F, Frydman SM, Monastyrskyi A, Roush WR, et al. Therapeutic Targeting of CDK12/CDK13 in Triple-Negative Breast Cancer. Cancer Cell 2019; 36(5):545–58 e7.

50. Hamilton MJ, Suri M. CDK13-related disorder. Adv Genet 2019; 103(163–82.

51. Rouxel F, Relator R, Kerkhof J, McConkey H, Levy M, Dias P, et al. CDK13-related disorder: Report of a series of 18 previously unpublished individuals and description of an epigenetic signature. Genet Med 2022; 24(5):1096–107.

52. Contro G, Baroni MC, Caraffi SG, Napoli M, Artuso R, Giliberti A, et al. CDK13-Related Disorder: Novel Insights From A Series of 27 Cases and Recommendations for Clinical Management. Clin Genet 2025; 108(2):146–55.

53. Fan Z, Devlin JR, Hogg SJ, Doyle MA, Harrison PF, Todorovski I, et al. CDK13 cooperates with CDK12 to control global RNA polymerase II processivity. Sci Adv 2020; 6(18).

54. Insco ML, Abraham BJ, Dubbury SJ, Kaltheuner IH, Dust S, Wu C, et al. Oncogenic CDK13 mutations impede nuclear RNA surveillance. Science 2023; 380(6642):eabn7625.

55. Zhang Y, Chen XN, Zhang H, Wen JK, Gao HT, Shi B, et al. CDK13 promotes lipid deposition and prostate cancer progression by stimulating NSUN5-mediated m5C modification of ACC1 mRNA. Cell Death Differ 2023; 30(12):2462–76.

56. Wu C, Xie T, Guo Y, Wang D, Qiu M, Han R, et al. CDK13 phosphorylates the translation machinery and promotes tumorigenic protein synthesis. Oncogene 2023; 42(16):1321–30.

57. Cheng L, Zhou S, Zhou S, Shi K, Cheng Y, Cai MC, et al. Dual Inhibition of CDK12/CDK13 Targets Both Tumor and Immune Cells in Ovarian Cancer. Cancer Res 2022; 82(19):3588–602.

58. Zeng Y, Wang S, Gao S, Soares F, Ahmed M, Guo H, et al. Refined RIP-seq protocol for epitranscriptome analysis with low input materials. PLoS Biol 2018; 16(9):e2006092.

59. Dong X, Chen G, Cai Z, Li Z, Qiu L, Xu H, et al. CDK13 RNA Over-Editing Mediated by ADAR1 Associates with Poor Prognosis of Hepatocellular Carcinoma Patients. Cell Physiol Biochem 2018; 47(6):2602–12.

60. Ramirez-Moya J, Miliotis C, Baker AR, Gregory RI, Slack FJ, Santisteban P. An ADAR1-dependent RNA editing event in the cyclin-dependent kinase CDK13 promotes thyroid cancer hallmarks. Mol Cancer 2021; 20(1):115.

61. Ishizuka JJ, Manguso RT, Cheruiyot CK, Bi K, Panda A, Iracheta-Vellve A, et al. Loss of ADAR1 in tumours overcomes resistance to immune checkpoint blockade. Nature 2019; 565(7737):43–8.

62. Jiang G, Wang Z, Cheng Z, Wang W, Lu S, Zhang Z, et al. The integrated molecular and histological analysis defines subtypes of esophageal squamous cell carcinoma. Nat Commun 2024; 15(1):8988.

63. Simon Andrews F, Segonds-Pichon A, Biggins L, Krueger C, Wingett S: FastQC: a quality control tool for high throughput sequence data. 2010.

64. Cheng AP, Huang L, Oberkofler L, Johnson NR, Glodeanu AS, Stillman K, et al. Fungal Argonaute proteins act in bidirectional cross-kingdom RNA interference during plant infection. Proc Natl Acad Sci U S A 2025; 122(17):e2422756122.

65. Frankish A, Carbonell-Sala S, Diekhans M, Jungreis I, Loveland JE, Mudge JM, et al. GENCODE: reference annotation for the human and mouse genomes in 2023. Nucleic Acids Res 2023; 51(D1):D942–D9.

66. Dobin A, Davis CA, Schlesinger F, Drenkow J, Zaleski C, Jha S, et al. STAR: ultrafast universal RNA-seq aligner. Bioinformatics 2013; 29(1):15–21.

67. Li B, Dewey CN. RSEM: accurate transcript quantification from RNA-Seq data with or without a reference genome. BMC Bioinformatics 2011; 12(323.

68. Ison J, Rapacki K, Menager H, Kalas M, Rydza E, Chmura P, et al. Tools and data services registry: a community effort to document bioinformatics resources. Nucleic Acids Res 2016; 44(D1):D38–47.

69. Van der Auwera GA, O’Connor BD: Genomics in the cloud: using Docker, GATK, and WDL in Terra. O’Reilly Media; 2020.

70. Danecek P, Bonfield JK, Liddle J, Marshall J, Ohan V, Pollard MO, et al. Twelve years of SAMtools and BCFtools. Gigascience 2021; 10(2).

71. Wang K, Li M, Hakonarson H. ANNOVAR: functional annotation of genetic variants from high-throughput sequencing data. Nucleic Acids Res 2010; 38(16):e164.

72. Schubach M, Maass T, Nazaretyan L, Roner S, Kircher M. CADD v1.7: using protein language models, regulatory CNNs and other nucleotide-level scores to improve genome-wide variant predictions. Nucleic Acids Res 2024; 52(D1):D1143–D54.

73. Stephenson JD, Totoo P, Burke DF, Janes J, Beltrao P, Martin MJ. ProtVar: mapping and contextualizing human missense variation. Nucleic Acids Res 2024; 52(W1):W140–W7.

74. c GPCCaAAaagcbAGRgue, 14 PgBCoMGRABEDHHYKVKCLSMD, MIT BIo, 13 HLESADMGSBGN, 31 CIfMRGNTLHGNPRAM, 5 IBDRGRHSJTKZ, et al. A global reference for human genetic variation. Nature 2015; 526(7571):68–74.

75. Sherry ST, Ward M-H, Kholodov M, Baker J, Phan L, Smigielski EM, et al. dbSNP: the NCBI database of genetic variation. Nucleic acids research 2001; 29(1):308–11.

76. Fu W, O’connor TD, Jun G, Kang HM, Abecasis G, Leal SM, et al. Analysis of 6,515 exomes reveals the recent origin of most human protein-coding variants. Nature 2013; 493(7431):216–20.

77. D’Addabbo P, Cohen-Fultheim R, Twersky I, Fonzino A, Silvestris DA, Prakash A, et al. REDIportal: toward an integrated view of the A-to-I editing. Nucleic Acids Res 2025; 53(D1):D233–D42.

78. Ritchie ME, Phipson B, Wu D, Hu Y, Law CW, Shi W, et al. limma powers differential expression analyses for RNA-sequencing and microarray studies. Nucleic Acids Res 2015; 43(7):e47.

79. Yu G, Wang LG, Han Y, He QY. clusterProfiler: an R package for comparing biological themes among gene clusters. OMICS 2012; 16(5):284–7.

80. Yu G, He QY. ReactomePA: an R/Bioconductor package for reactome pathway analysis and visualization. Mol Biosyst 2016; 12(2):477–9.

81. Love MI, Huber W, Anders S. Moderated estimation of fold change and dispersion for RNA-seq data with DESeq2. Genome Biol 2014; 15(12):550.

82. Zhang Y, Liu T, Meyer CA, Eeckhoute J, Johnson DS, Bernstein BE, et al. Model-based analysis of ChIP-Seq (MACS). Genome Biol 2008; 9(9):R137.

83. Heinz S, Benner C, Spann N, Bertolino E, Lin YC, Laslo P, et al. Simple combinations of lineage-determining transcription factors prime cis-regulatory elements required for macrophage and B cell identities. Molecular cell 2010; 38(4):576–89.

84. Hofacker IL, Priwitzer B, Stadler PF. Prediction of locally stable RNA secondary structures for genome-wide surveys. Bioinformatics 2004; 20(2):186–90.

85. Reuter JS, Mathews DH. RNAstructure: software for RNA secondary structure prediction and analysis. BMC Bioinformatics 2010; 11(129.

86. Stellos K, Gatsiou A, Stamatelopoulos K, Perisic Matic L, John D, Lunella FF, et al. Adenosine-to-inosine RNA editing controls cathepsin S expression in atherosclerosis by enabling HuR-mediated post-transcriptional regulation. Nat Med 2016; 22(10):1140–50.

87. Qi JC, Yang Z, Lin T, Ma L, Wang YX, Zhang Y, et al. CDK13 upregulation-induced formation of the positive feedback loop among circCDK13, miR-212-5p/miR-449a and E2F5 contributes to prostate carcinogenesis. J Exp Clin Cancer Res 2021; 40(1):2.

