## Supplementary Figures for "The RNA Editome of Esophageal Squamous Cell Carcinoma Identifies an ADAR1-CDK13 Editing Feedback Loop Mediating cGAS–STING Activation in Early Tumorigenesis"

**a**

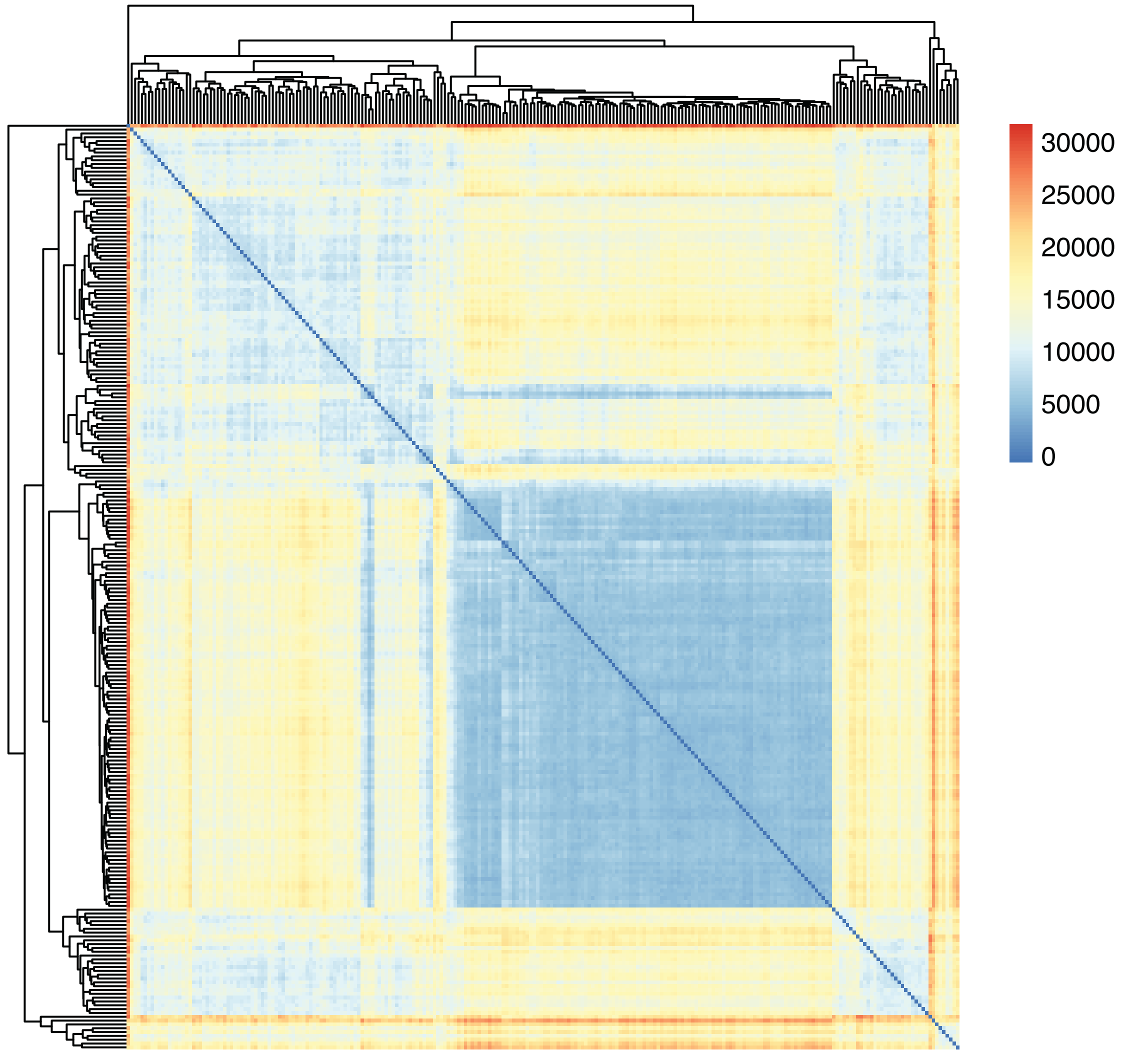


**b**

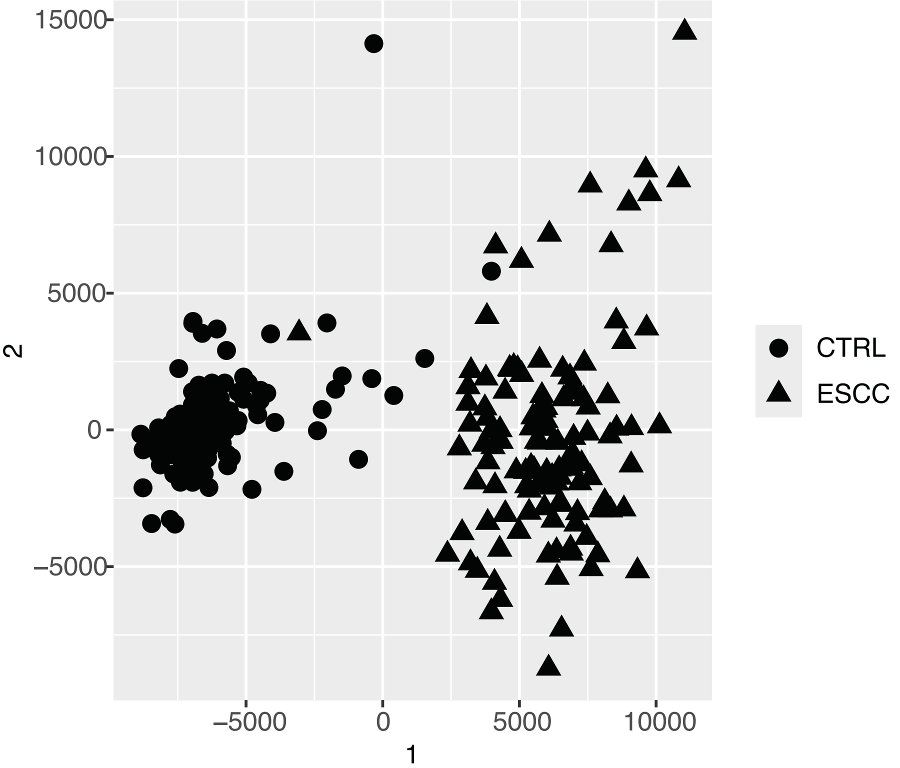


**Supplementary Fig. 1. Cluster of ESCC and matched adjacent normal tissues. (a)** Heatmap showed the clustering of samples. Poisson distance was measured for clustering of 121 tumor and matched normal adjacent samples. **(b)** Multidimensional scaling plot showed clustering across samples.

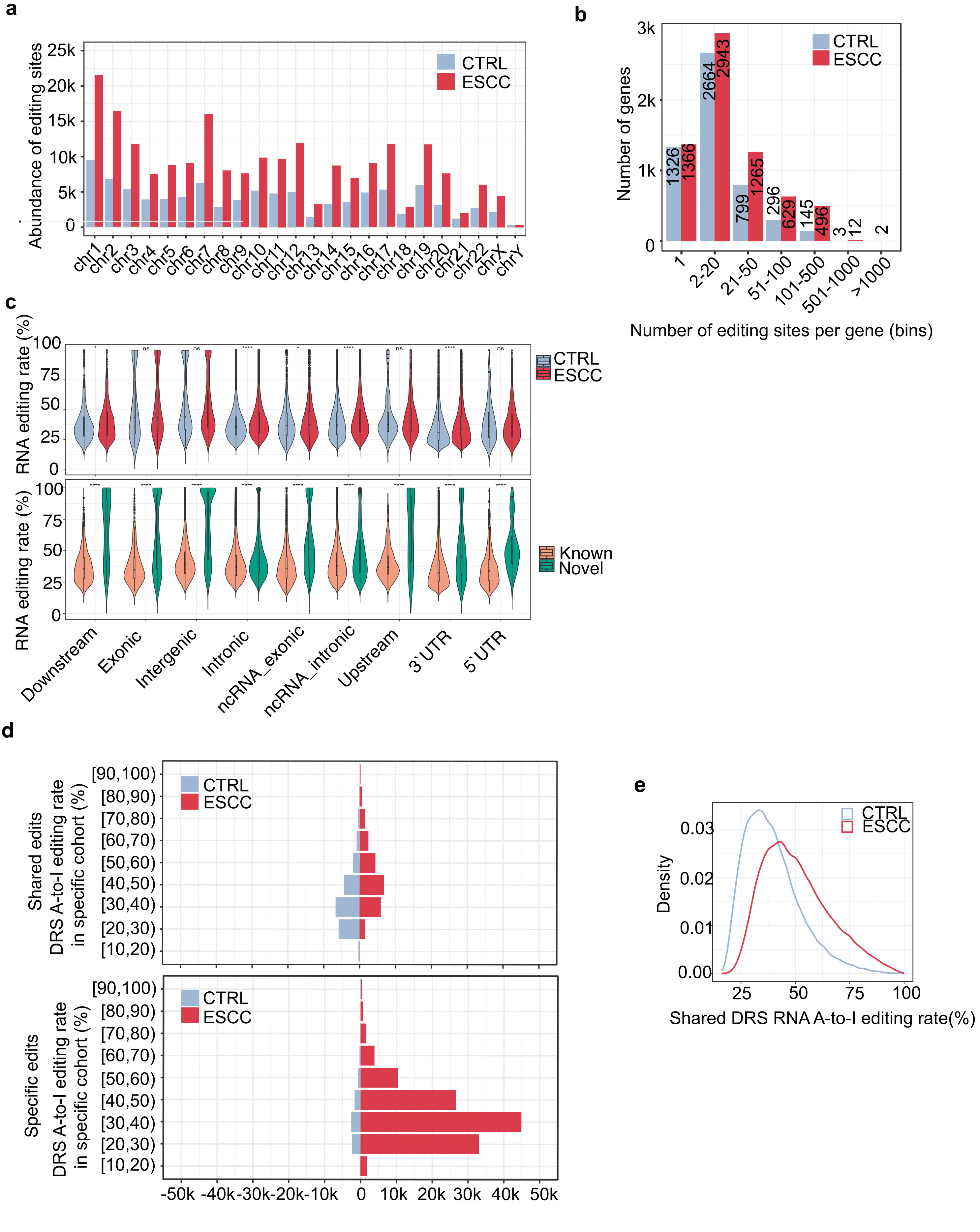


**Supplementary Fig. 2. RNA editing sites across samples.** **(a)** Distribution of the abundance for RNA editing sites across chromosomes. **(b)** Genes containing a binned number of editing sites in ESCC and CTRL samples. **(c)** Genomic loci of variants with RNA editing rate along chromosomes in ESCC and CTRL tissues or in known and novel edits. Significance was assessed by the Wilcoxon test and asterisks indicate significance with *, **, ***, and **** representing $p < 0.05$, $p < 0.01$, $p < 0.001$, $p < 0.0001$, respectively. **(d)** Comparison of editing rate for shared and ESCC/CTRL-specific A-to-I DRS edits and corresponding counts in each cohort. **(e)** Density of shared A-to-I DRS editing rate in ESCC and CTRL cohorts.

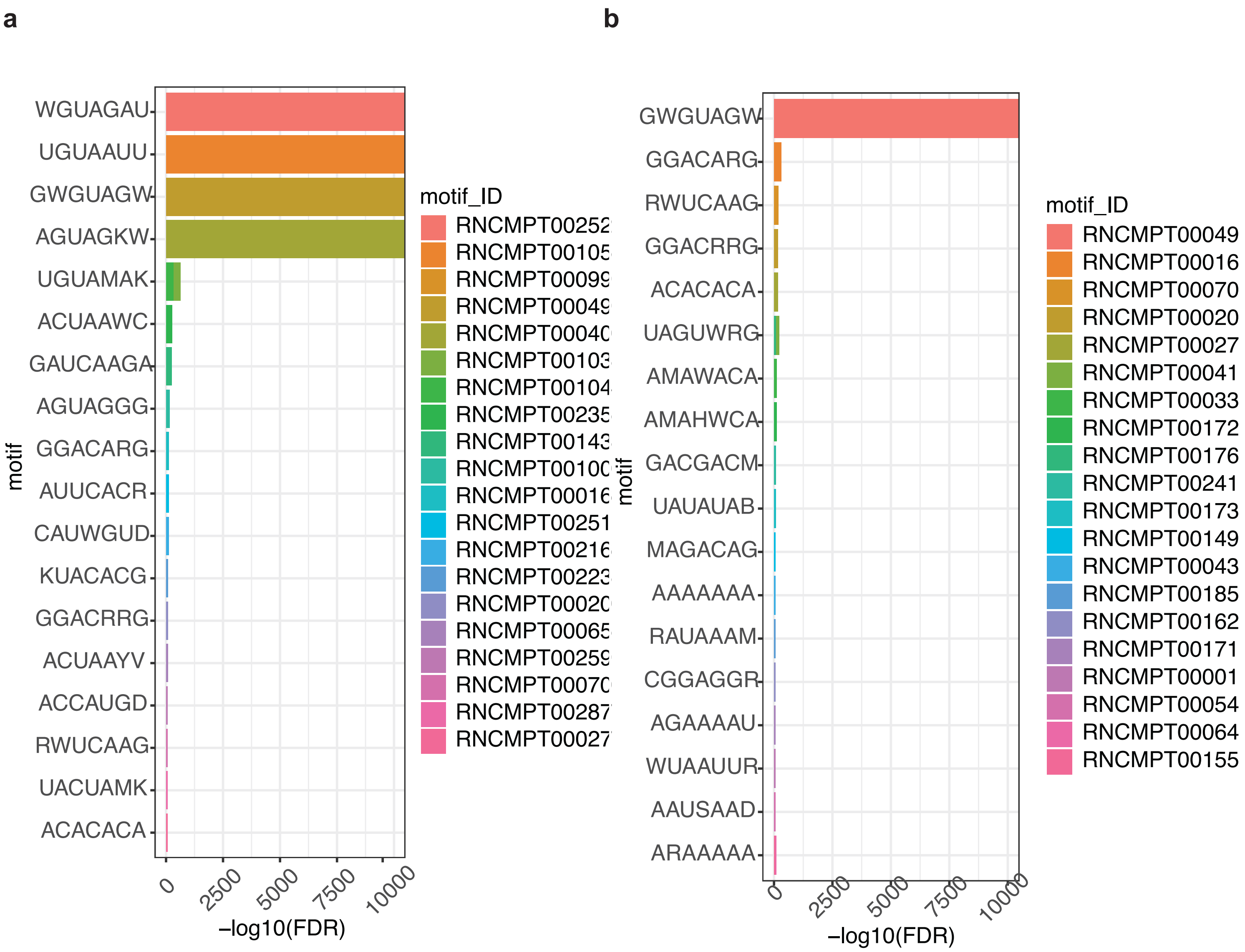


**Supplementary Fig. 3. Sequence motif enrichment analysis. (a, b)** 8-bp window surrounding DRS (±4 bp) in CTRL (a) and ESCC (b) were retained for analysis on Analysis of Motif Enrichment (AME). Default shuffled sequences were used as control. Average odds score was chose as scoring method, and fisher's exact test was selected for enrichment calculation.

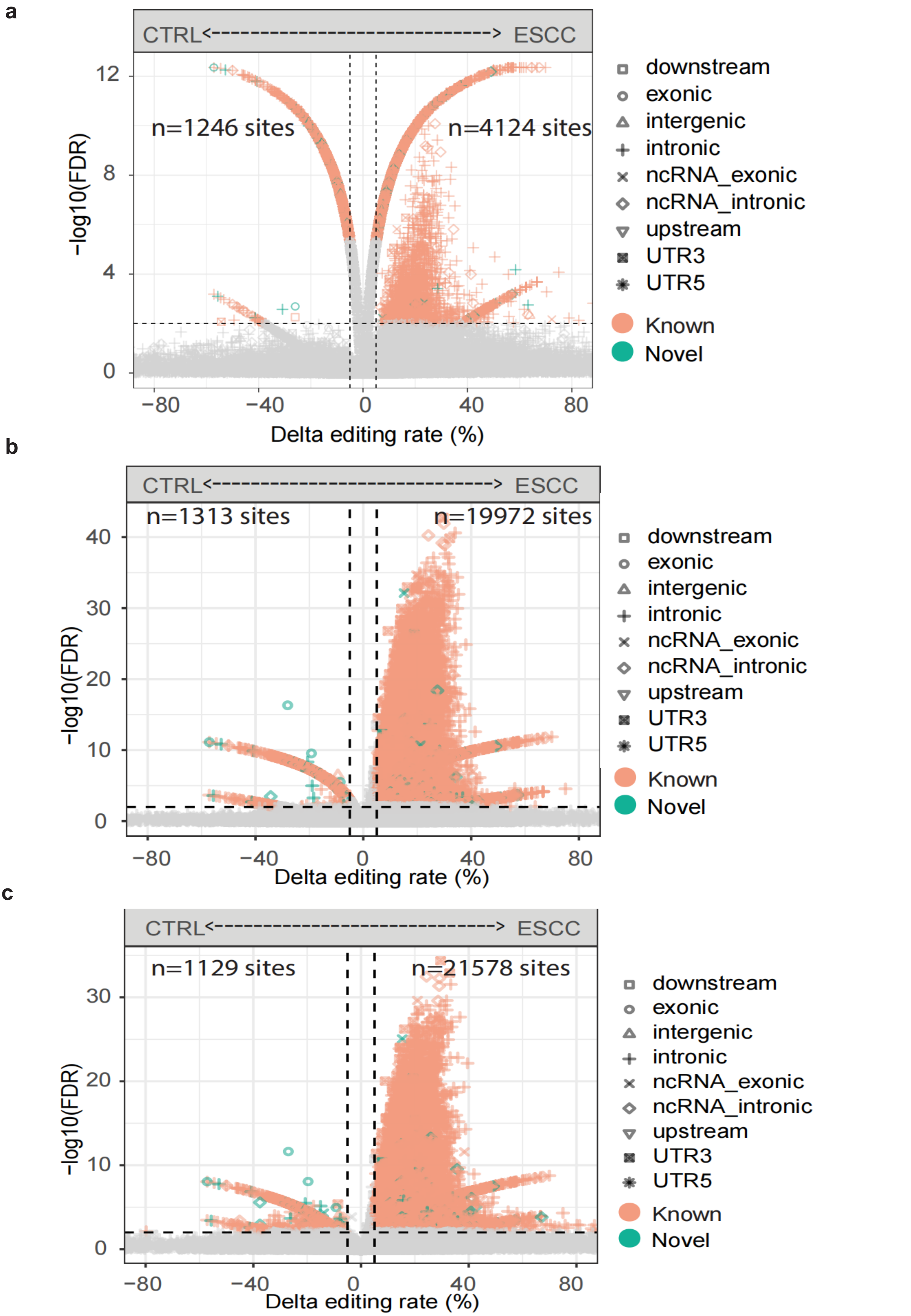


**Supplementary Fig. 4. Volcano plot presenting shared differentially edited sites (DRS) in ESCC and CTRL samples. (a-c)** Expression of *ADAR*1 **(a)**, *ADAR2* **(b)**, or *ADAR3* **(c)** was covariated indifferentia analysis, respectively. Shapes represent distinct variant types in ESCC and CTRL samples. Orangeshapes refer to known editing sites recorded in REDIportal and green shapes refer to identified novel editing sites.

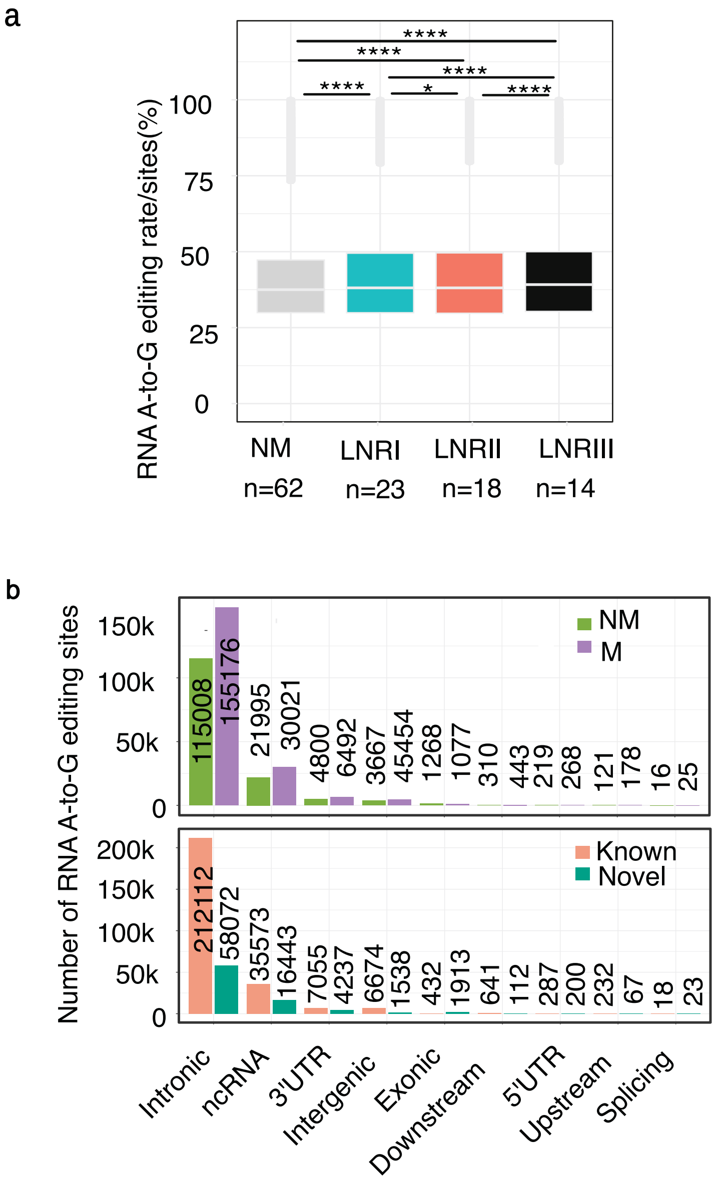


**Supplementary Fig. 5. Comparison of RNA A-to-I editing rates/sites in non-metastatic and distinct stages of metastatic ESCC. (a)** Metastatic ESCC were categorized to LNRI (n = 23), LNRII (n = 18), and LNRIII (n = 14) based on the status of lymph node. Asterisks *, **, *** and **** indicated significance representing $p < 0.05$, $p < 0.01$, $p < 0.001$, $p < 0.0001$, respectively. **(b)** Variant types for RNA A-to-I edits in NM and M samples, and A-to-I edits that were classified as known or novel.

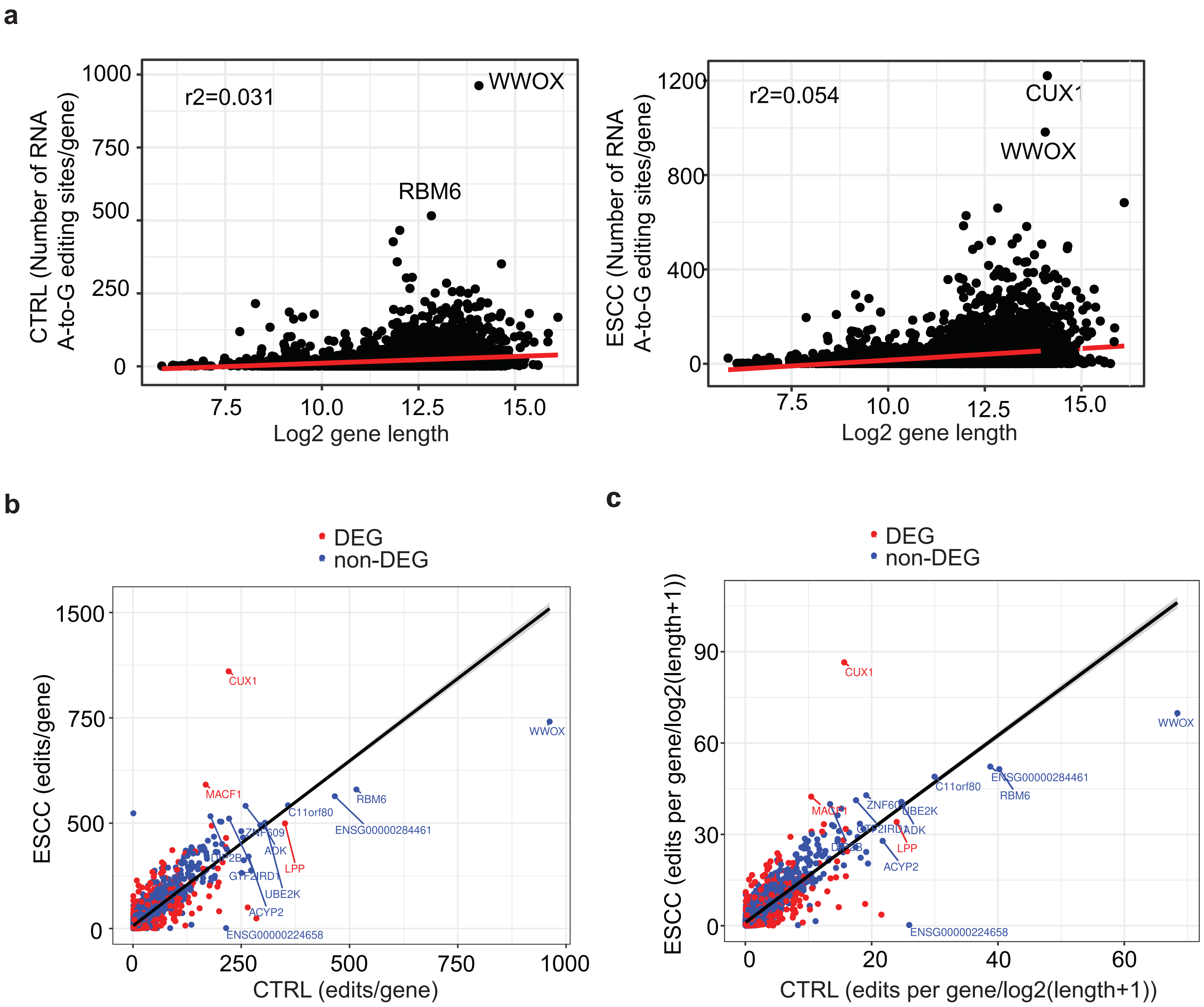


**Supplementary Fig. 6. Association analysis of gene length with RNA editing sites. (a)** Linear regression was conducted to measure the relationship of gene length and counts of RNA A-to-I editing sites across genes in ESCC and CTRL samples. **(b)** Pairwise comparisons of concordance for the abundance of RNA editing sites across genes in ESCC and CTRL samples. Genes differentially expressed were labeled in red and non-differentially expressed were in blue. **(c)** Pairwise comparisons of concordance for the abundance of RNA editing sites normalized by gene length across genes in ESCC and CTRL samples. Genes differentially expressed were labeled in red and non-differentially expressed were in blue.

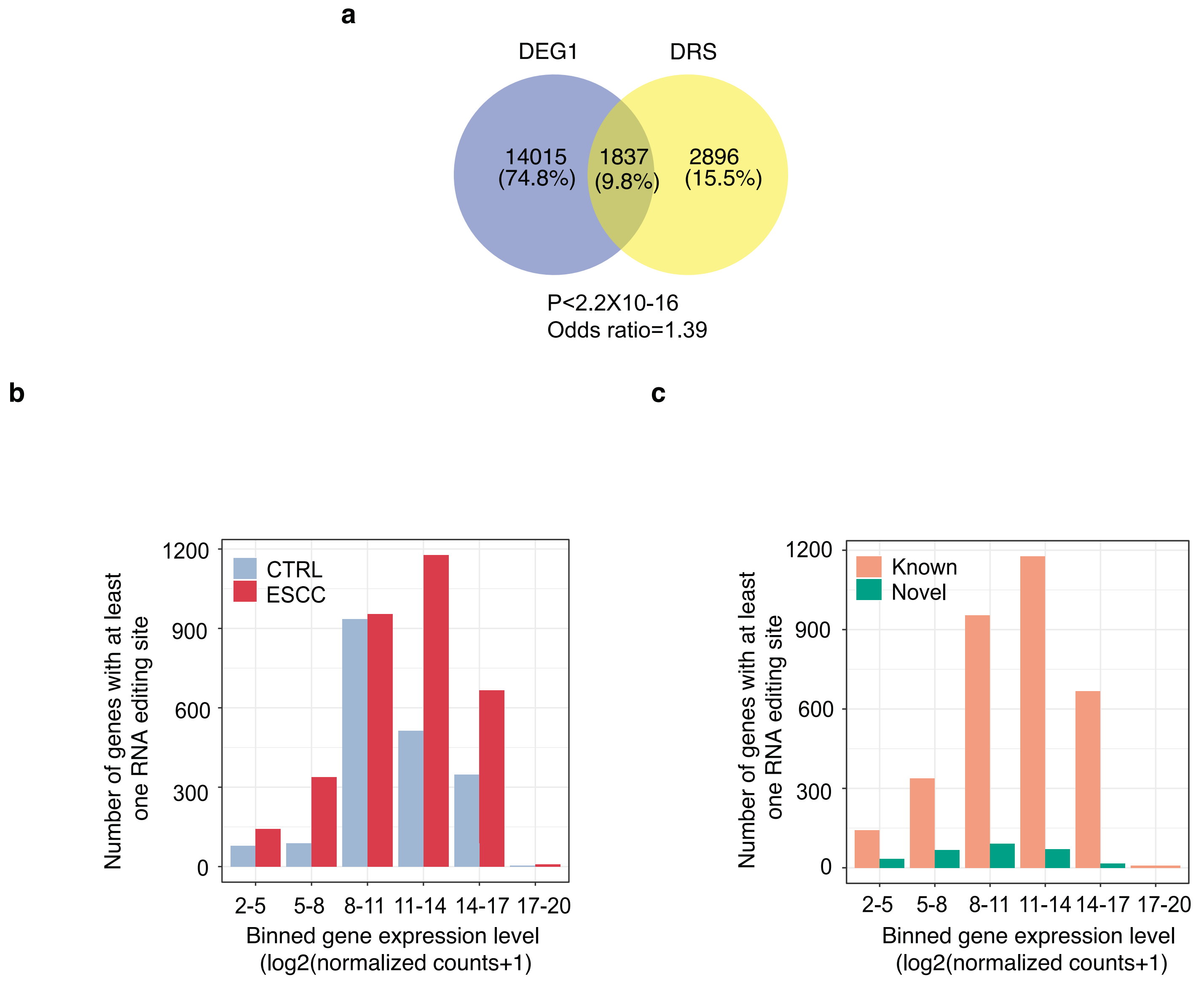


**Supplementary Fig. 7. Distribution of RNA-edited genes across expression bins. (a)** Venn diagram of DEG1 and DRS identified from ESCC and CTRL samples. **(b)** Counts of genes containing at least one RNA editing site binned by expression level in ESCC and CTRL samples. **(c)** Counts of genes containing at least one RNA editing site classified as known or novel across expression bins.

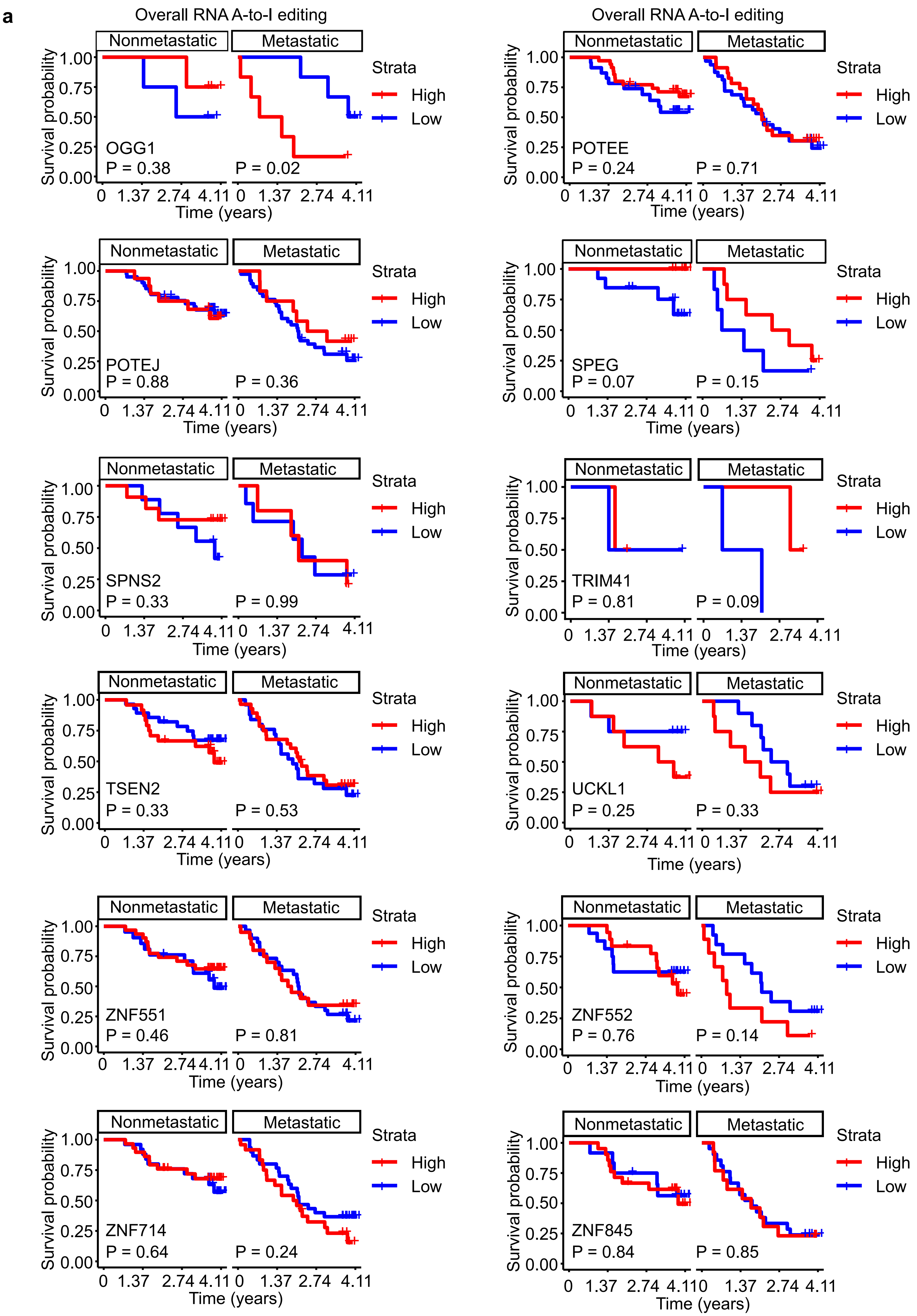
a
Overall RNA A-to-I editing
Overall RNA A-to-I editing
Nonmetastatic
Metastatic
Strata
High
Low
Survival probability
Time (years)
1.00
0.75
0.50
0.25
0.00
0
1.37
2.74
4.11
OGG1
P = 0.38
P = 0.02
POTEE
P = 0.24
P = 0.71
POTEJ
P = 0.88
P = 0.36
SPEG
P = 0.07
P = 0.15
SPNS2
P = 0.33
P = 0.99
TRIM41
P = 0.81
P = 0.09
TSEN2
P = 0.33
P = 0.53
UCKL1
P = 0.25
P = 0.33
ZNF551
P = 0.46
P = 0.81
ZNF552
P = 0.76
P = 0.14
ZNF714
P = 0.64
P = 0.24
ZNF845
P = 0.84
P = 0.85

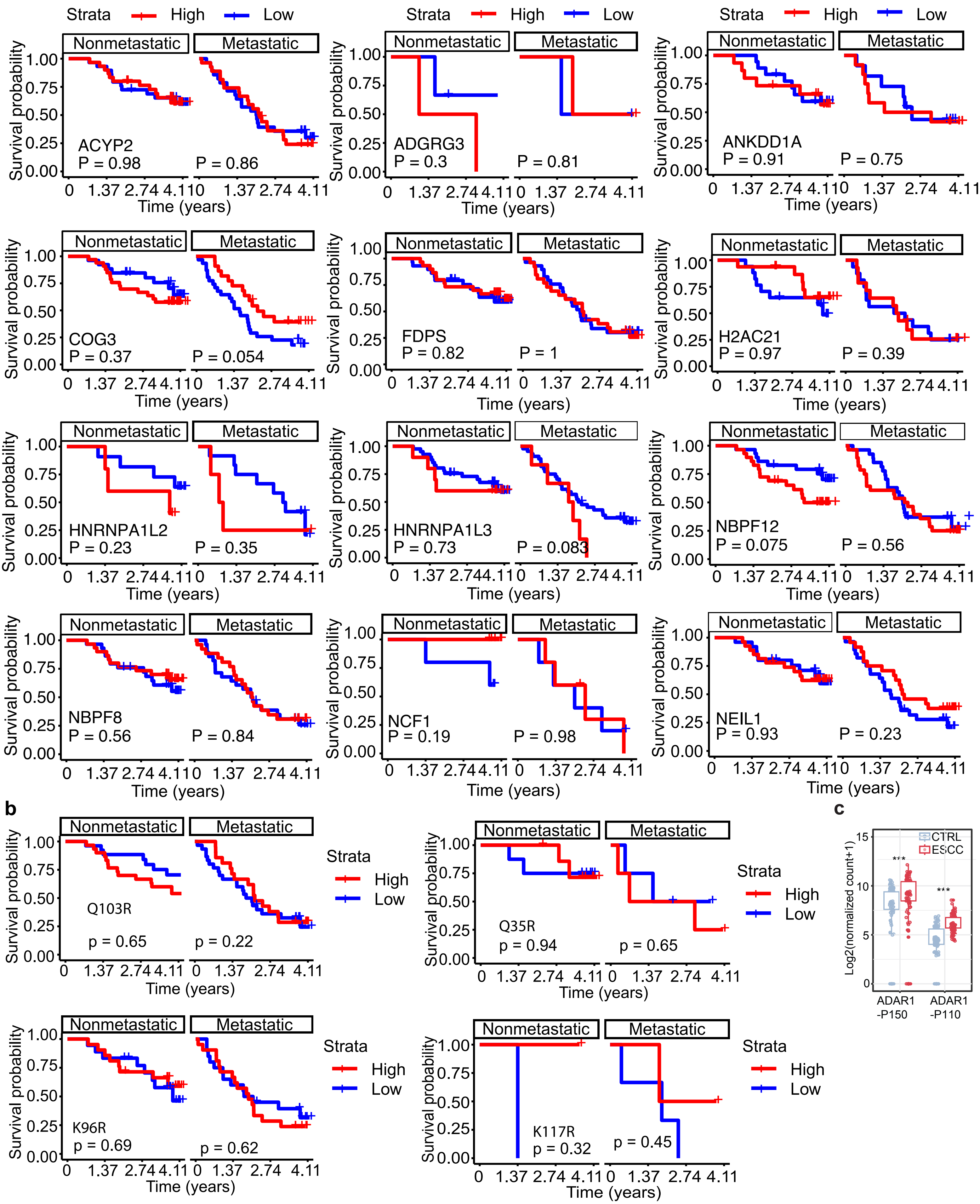


**Supplementary Fig. S8. Survival analysis of overall RNA A-to-I editing. (a,b)** Kaplan-Meier curves for patiens stratified by ESCC metastasis. Survival of overall RNA A-to-I editing in 24 RNAs **(a)** and 4 recoding sites **(b)**. Kaplan-meier analysis was stratified by non-metastatic and metastatic ESCC. RNA editing levels were divided into high (red line) and low (blue line) groups based on their median values. NM: nonmetastatic, M: metastatic.

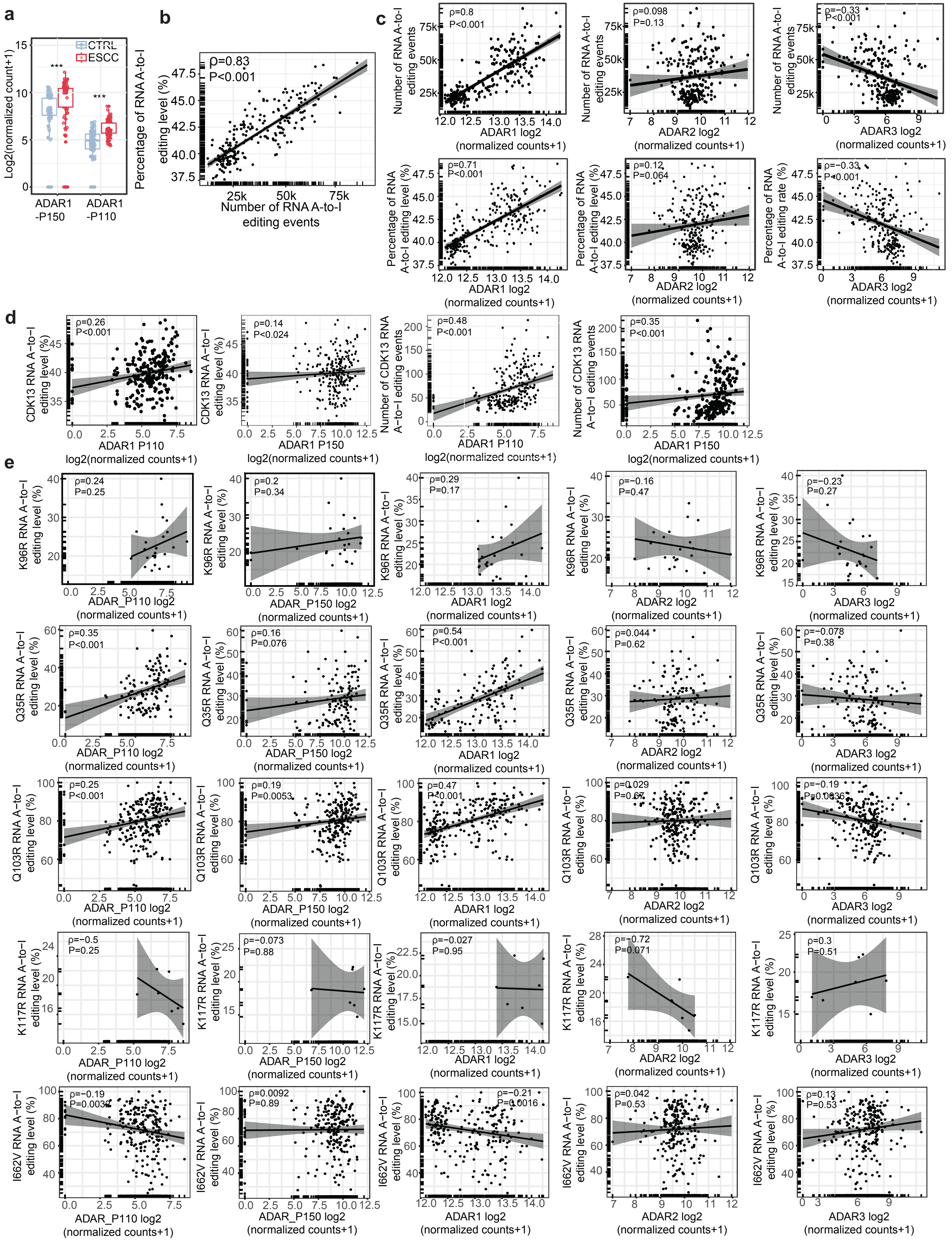

**Supplementary Fig. S9. Association of ADAR expression with RNA diting level. (a)** The expression of ADAR1 P150 and P110 in ESCC and CTRL. **(b)** Correlation of RNA editing level wih RNA editing events. **(c)** Correlation of RNA editing level or events with ADAR expression. **(d)** Correlation of *CDK13* RNA editing level or events with ADAR1 P150 and P110 expression. **(e)** Correlation of RNA editing level in 5 recod-ing sites in *CDK13* with ADAR expression.

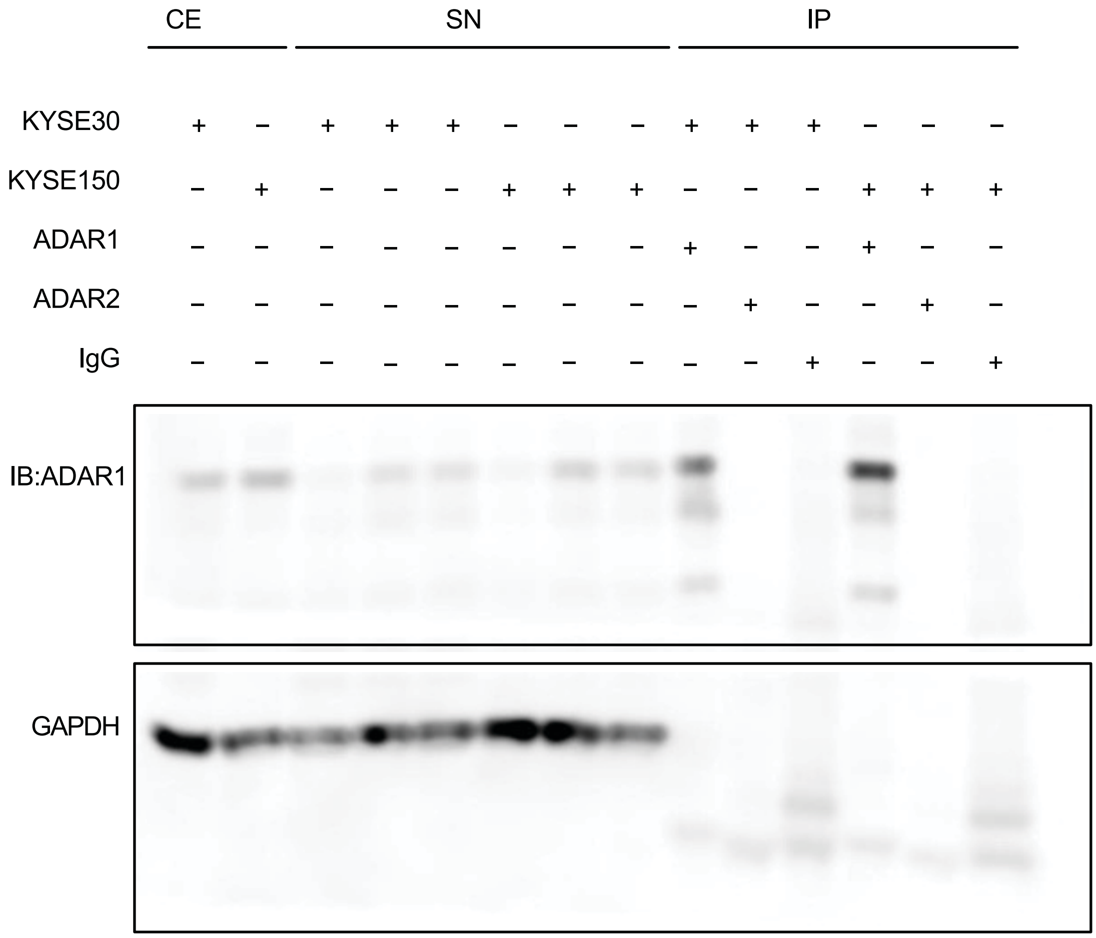


**Supplementary Fig. S10. ADAR1 RNA immunoprecipitation.** Immunoblotting examining ADAR1 abundance in distinct fractions of ADAR1 immunoprecipitation. GAPDH was used as loading control. Abbreviations represent CE: crude extract; SN: supernatant; IP: Immunoprecipitation.

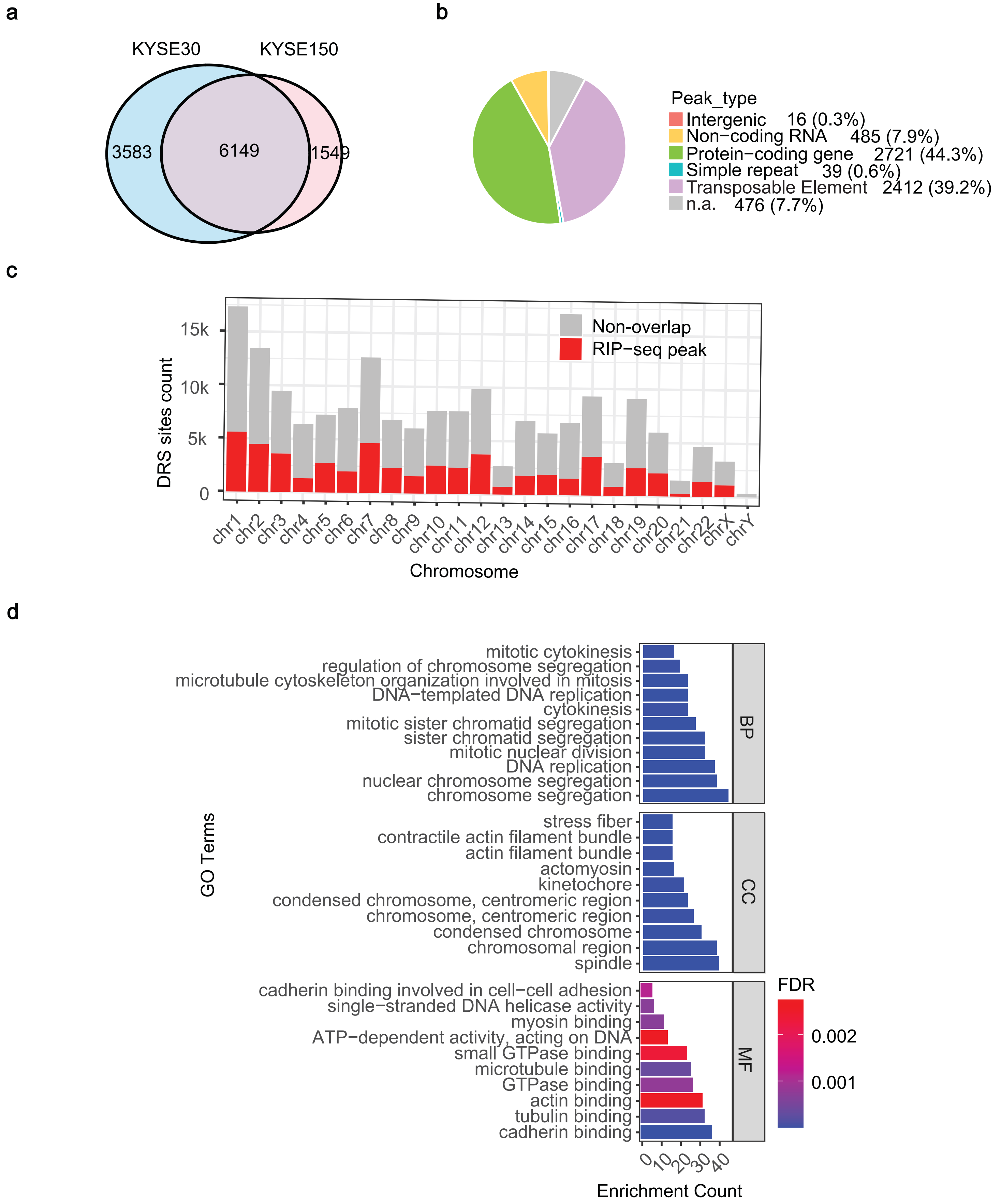


**Supplementary Fig. 11. RIP-seq analysis revealed ADAR1 associated peaks. (a)** Venn diagram presenting detected peaks in KYSE30 and KYSE150. **(b)** Pie chart presenting distribution of overlapping peaks across genome. **(c)** Overlay bar chart presenting counts of DRS across chromosomes. **(d)** Top 10 GO terms of BP, CC and MF enriched among ADAR1-dependent DRS in genes. DRS: differential RNA editing sites; BP: biological process; CC: cellular component; MF: molecular function.

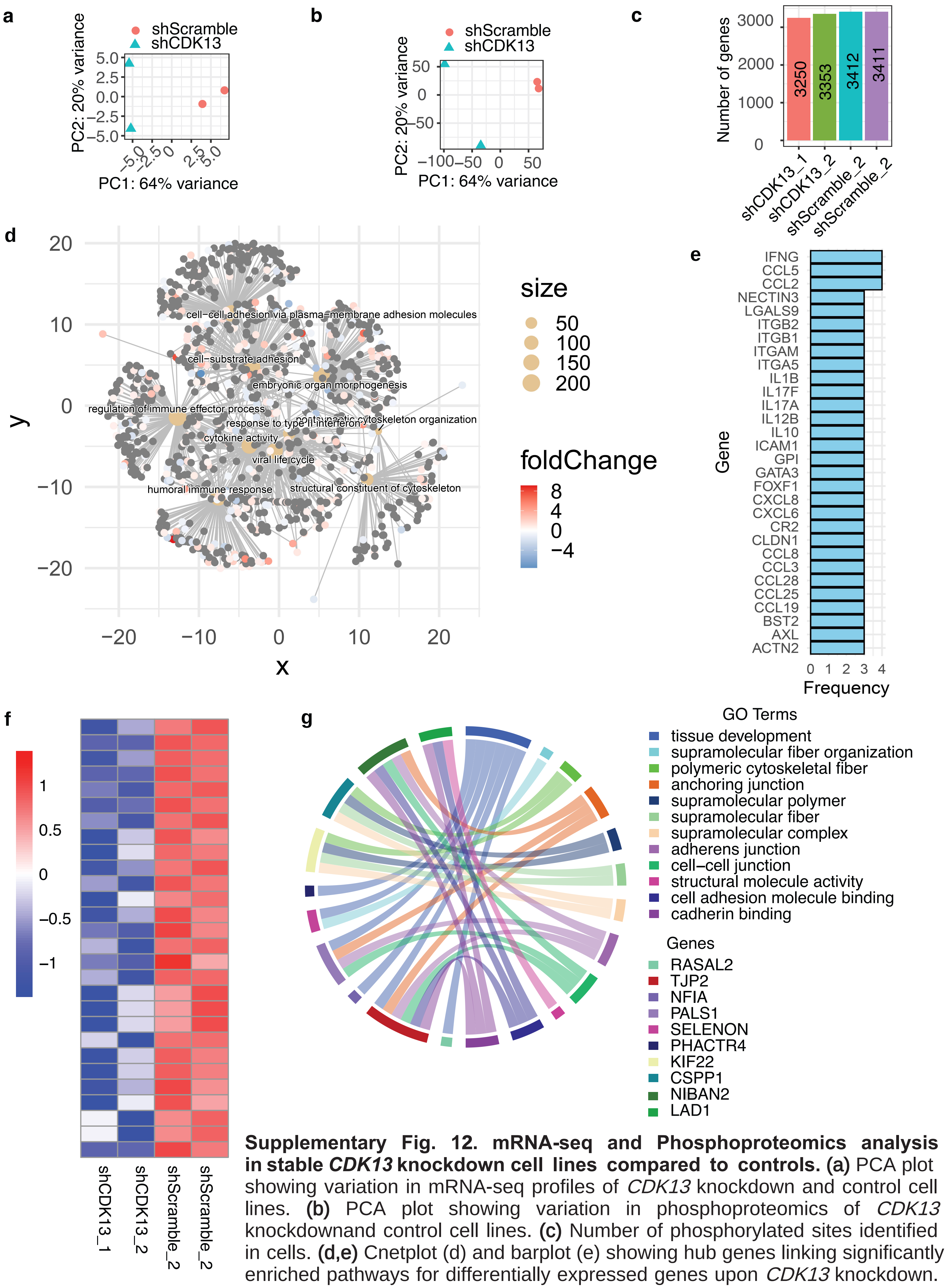


**Supplementary Fig. 12. mRNA-seq and Phosphoproteomics analysis in stable *CDK13* knockdown cell lines compared to controls. (a)** PCA plot showing variation in mRNA-seq profiles of *CDK13* knockdown and control cell lines. **(b)** PCA plot showing variation in phosphoproteomics of *CDK13* knockdownand control cell lines. **(c)** Number of phosphorylated sites identified in cells. **(d,e)** Cnetplot (d) and barplot (e) showing hub genes linking significantly enriched pathways for differentially expressed genes upon *CDK13* knockdown. **(f)** Heatmap showing phosphorylation for 28 candidate sites of CDK13. **(g)** ChordDiagram showing GO terms enriched among 22 proteins containing potential phosphorylation sites of CDK13.

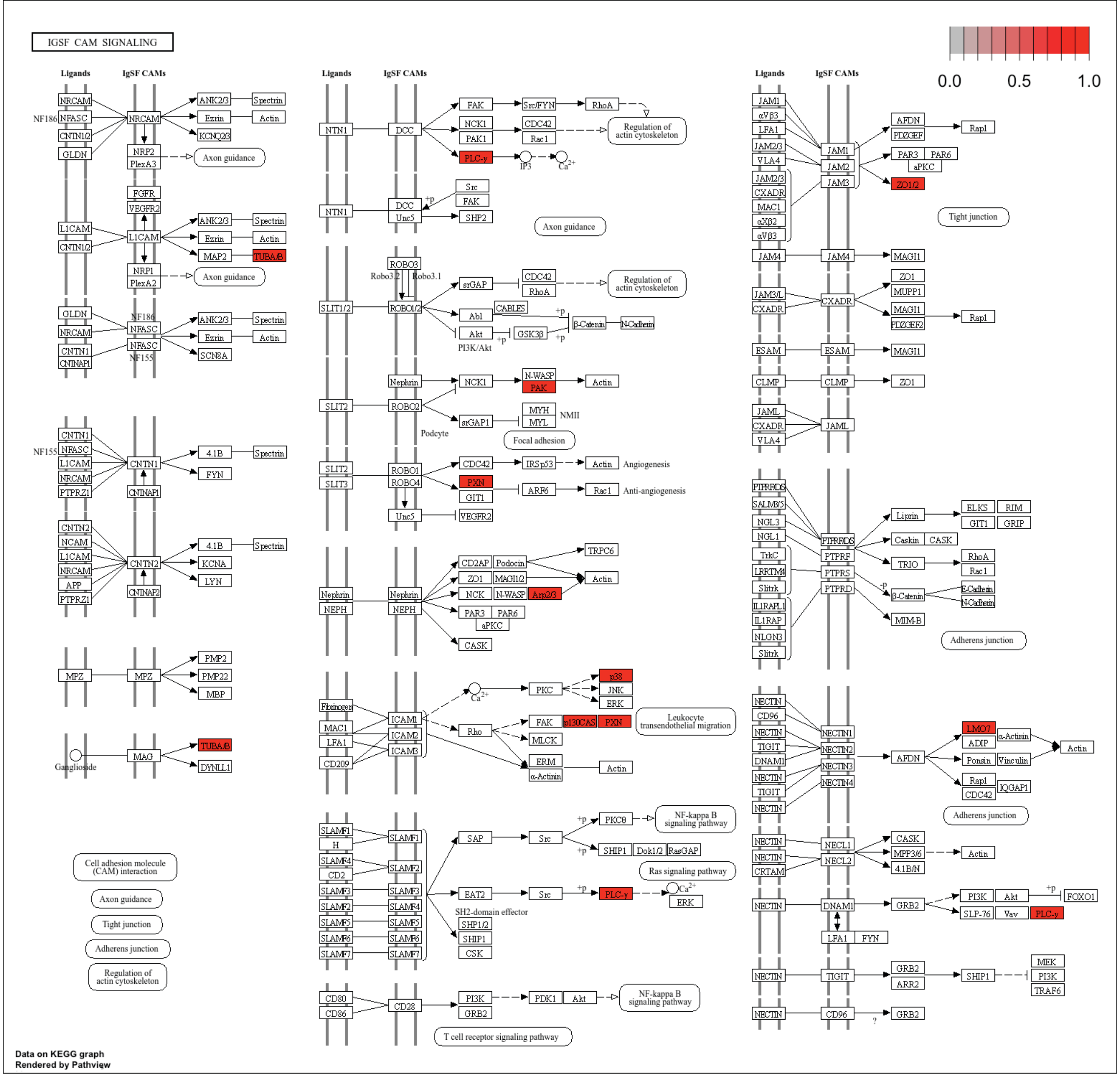


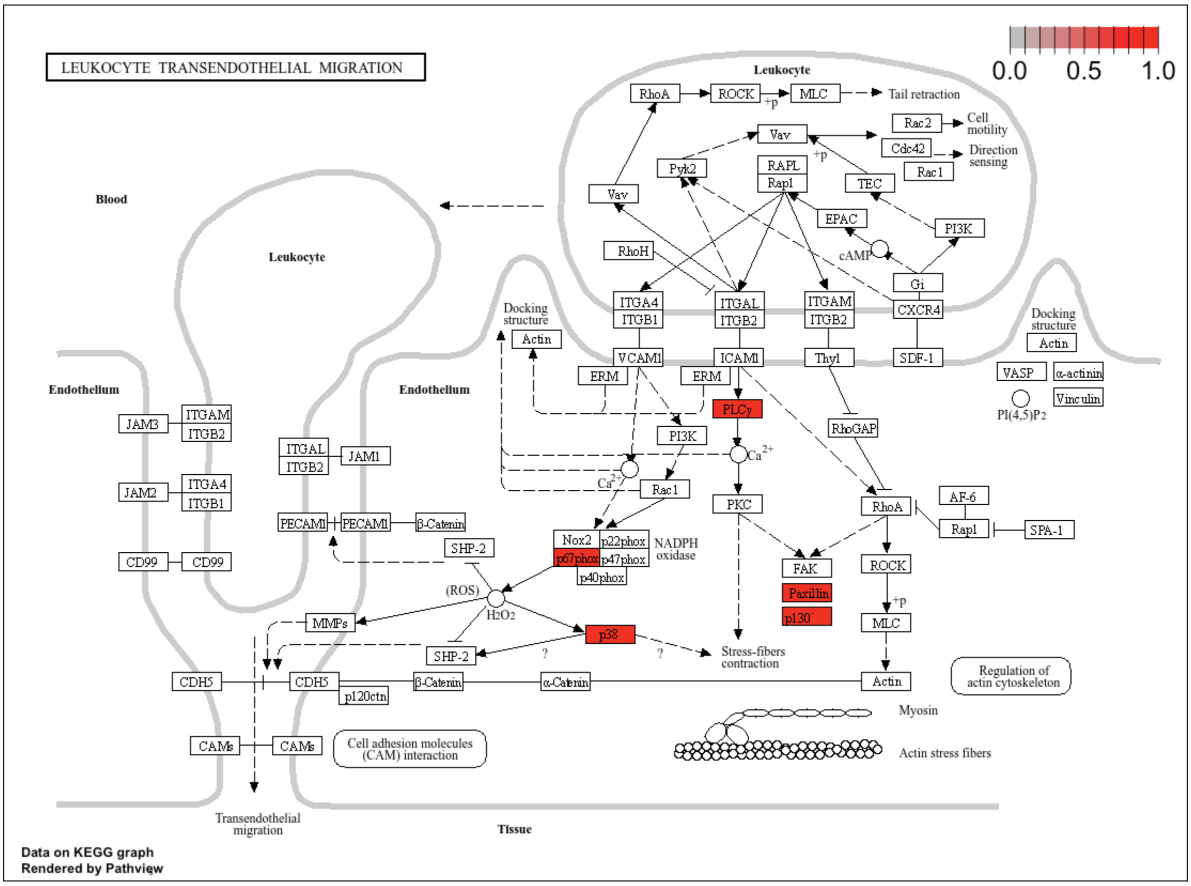


**Supplementary Fig. 13. *CDK13* editing in ESCC may regulate IgSF CAM signalling and Leukocyte transendothelial migration.** IgSF CAM signalling and Leukocyte transendothelial migration were enriched among 59 genes, which involved in 4 genes (encoding TUBA1B, ARPC2, PAK2, TJP2) in IgSF CAM signalling and 1 genes (encoding NCF2) in Leukocyte transendothelial migration, and 4 genes (encoding PLCG1, PXN, MAK13, BCAR1) in both pathways.
